# Queryome: Orchestrating Retrieval, Reasoning, and Synthesis across Biomedical Literature

**DOI:** 10.64898/2025.12.22.696019

**Authors:** Pranav Punuru, Nabil Ibtehaz, Swagarika Giri, Harsha Srirangam, Emilia Tugolukova, Daisuke Kihara

## Abstract

The rapid expansion of biomedical literature has made comprehensive manual synthesis increasingly difficult to perform effectively, creating a pressing need for AI systems capable of reasoning across verified evidence rather than merely retrieving it. However, existing retrieval-augmented generation (RAG) methods often fall short when faced with complex biomedical questions that require iterative reasoning and multi-step synthesis. Here, we developed Queryome, a deep research system consisting of specialized large language model (LLM) agents that can adapt their orchestration dynamically to a wide range of queries. Using a hybrid semantic–lexical retrieval engine spanning 28.3 million PubMed abstracts, it performs iterative, evidence-grounded synthesis. On the MIRAGE benchmark, Queryome achieved 88.98 % accuracy, surpassing prior systems by up to 14 points, and improved reasoning accuracy on the biomedical Human’s Last Exam (HLE) subset from 15.8% to 19.3%. Moreover, in a task for constructing a review article, it earned the highest composite score in comparison with Deep Research from OpenAI, Google, Perplexity, and Scite.AI, reflecting its strong literature retrieval and synthesis capabilities.

## Introduction

The rapid expansion of biomedical literature presents a major challenge for researchers, clinicians, and scientists. The number of publications indexed in databases such as PubMed [1] has grown exponentially, making it practically impossible for individuals to remain current with all relevant findings in their field [2]. This deluge of information necessitates the development of advanced computational tools capable of not only retrieving but also synthesizing knowledge at scale.

Large Language Models (LLMs) have emerged as powerful tools for processing and generating natural language, offering the potential to automate aspects of literature review [3–5]. However, their application in biomedicine is constrained by several domain-specific limitations. The scarcity and fragmentation of high-quality biomedical training data present significant challenges, as medical data is frequently siloed across different institutions and stored in various formats [6]. Consequently, LLMs demonstrate a limited grasp of biomedical language [7]. General-domain LLMs often lack specialized medical knowledge because they are primarily trained on non-medical datasets, limiting their effectiveness in healthcare settings [8]. Beyond these domain-specific challenges, LLMs across all domains are susceptible to hallucination; a phenomenon where models generate plausible yet factually incorrect or unsubstantiated information [9]. These combined limitations of LLMs and their reliance on static, pre-trained knowledge that may not include the latest research findings restrict their utility in dynamic fields like biomedicine, where traceable evidence and up to date information is crucial.

To overcome the static knowledge and factual reliability issues inherent to standalone LLMs, researchers have explored methods to link model outputs with external sources of information. One such method is Retrieval-Augmented Generation (RAG) where LLM outputs are grounded in specific, verifiable information [10]. In a traditional RAG workflow, the LLM is assisted or complemented with external knowledge sources by first retrieving relevant documents from a specialized database based on the user’s query; these retrieved materials are then supplied as contextual evidence to guide the model’s response generation. For biology, recent RAG methods such as BiomedRAG [11] and BioRAG [12] have demonstrated a substantial improvement in question-answering performance. The authors of BiomedRAG report a 9.95% improvement on average on their dataset, and even outperform baseline LLMs by 4.97% [12]. However, conventional RAG systems applied to complex biomedical question-answering datasets such as MIRAGE [13] have shown limitations. These systems often struggle with the nuanced, multi-faceted nature of biomedical queries, indicating that simple retrieval and synthesis are insufficient for deep scientific inquiry.

More recently, the concept of agentic RAG has gained traction, promising more sophisticated "deep research" capabilities [14, 15]. Systems developed by industry leaders such as OpenAI [16], Perplexity AI [17], and Google [18] have demonstrated the ability to decompose complex questions, perform iterative searches, and synthesize more comprehensive reports. Yet, these general-purpose agentic systems are not specifically tailored for the biomedical domain. They may lack access to curated, comprehensive biomedical corpora and often perform a high-level synthesis without reasoning deeply over the full context of every single retrieved document, potentially missing critical details or conflicting evidence.

To bridge this gap, we introduce Queryome, a multi-agent deep research system designed specifically for end-to-end biomedical literature analysis. Queryome orchestrates a hierarchy of collaborating AI agents that perform iterative, multi-faceted searches against a curated, comprehensive search engine covering the entirety of PubMed [1]. Crucially, the system is engineered to reason over abstract text of every retrieved article, ensuring that its final synthesis is deeply grounded in the available evidence. By combining a specialized hybrid search system with a multi-agent reasoning architecture, Queryome aims to provide a robust, transparent, and powerful tool for navigating the vast biomedical literary landscape. Queryome demonstrates substantial improvements over existing biomedical RAG systems. On the MIRAGE benchmark [13], it achieves an average accuracy of 88.98 %, outperforming the best prior methods by up to 14 percentage points. On the biomedical subset of the Humanity’s Last Exam (HLE) [19], Queryome improves reasoning accuracy from 15.8% to 19.3%. In the Review Generation Test, it attains the top composite score (52 ± 1.6), with particularly high marks in Source Quality & Coverage (9.2 ± 0.3) and Formatting & Presentation (9.0 ± 0.4), demonstrating its superior retrieval precision and synthesis coherence.

## Methods

Queryome is an agentic RAG framework designed for biological knowledge retrieval, higher-order reasoning, and in-depth analysis of biomedical literature. Its curated RAG database is built from 28.3 million PubMed abstracts spanning diverse fields of biology and medicine. At the core of Queryome lies a reasoning-focused LLM, termed the Principal Investigator (PI) agent, which governs the overall research workflow. The PI orchestrates the process by delegating subtasks to specialized sub-agent teams, each consisting of a planner and a critic agent. The planner sub-agent strategically retrieves information from the database by integrating both semantic and lexical signals for comprehensive coverage, while the critic sub-agent evaluates the relevance and sufficiency of the retrieved evidence. Through iterative refinement, the sub-agents collaboratively generate detailed responses supported by evidence, which are then analyzed by the PI. The final output is compiled by a synthesizer agent into a coherent, citation-grounded summary. This architecture enables Queryome to scale its reasoning depth with query complexity while maintaining interpretability through explicit evidence tracking.

### Dataset Construction

We constructed a comprehensive biomedical literature database by processing the complete PubMed [1] Baseline collection distributed as XML files from the National Library of Medicine. The raw corpus initially contained approximately 40 million records (As of September 2025). Since abstract text is essential for both semantic indexing and retrieval quality, we applied a filtering criterion to retain only articles with available abstracts, excluding those with only titles and metadata. This filtering resulted in a final corpus of 28.3 million articles with complete abstracts. The filtered articles were parsed from XML and stored in a SQLite database containing fields for vector identifiers, PubMed identifiers, titles, abstracts, journal names, publication years, along with structured metadata fields for authors, MeSH terms, author keywords, and citations stored as JSON arrays. The vector identifier serves as the primary key and maps each embedding to its corresponding paper, enabling retrieval of full article metadata at inference time when the system identifies relevant documents.

Dense vector representations were computed for all 28.3 million articles to enable semantic retrieval. For each article, we concatenated the title and abstract into a single text passage for embedding generation. These passages were then embedded using the Linq-Embed-Mistral [20] model, a sentence transformer optimized for semantic similarity tasks. This model was selected due to its high performance on retrieval tasks, achieving a score of 60.2 on the Massive Text Embedding Benchmark (MTEB) leaderboard [21] as of May 29, 2024. The model generates 4,096-dimensional vectors for each article after which we assign the vector identifier to map the embeddings to the article’s information.

### Retrieval Architecture

The retrieval system combines dense semantic search with sparse lexical matching through a hybrid architecture (**Fig. 1a**). Dense retrieval employs FAISS (Facebook AI Similarity Search), a library optimized for efficient similarity search in high-dimensional vector spaces [22]. Given 28.3 million 4,096-dimensional embeddings, exact nearest-neighbor search would require computing distances to all vectors for each query, resulting in prohibitive computational costs. FAISS addresses this through approximate nearest-neighbor algorithms that trade minimal accuracy for substantial speed improvements.

**Figure 1.**
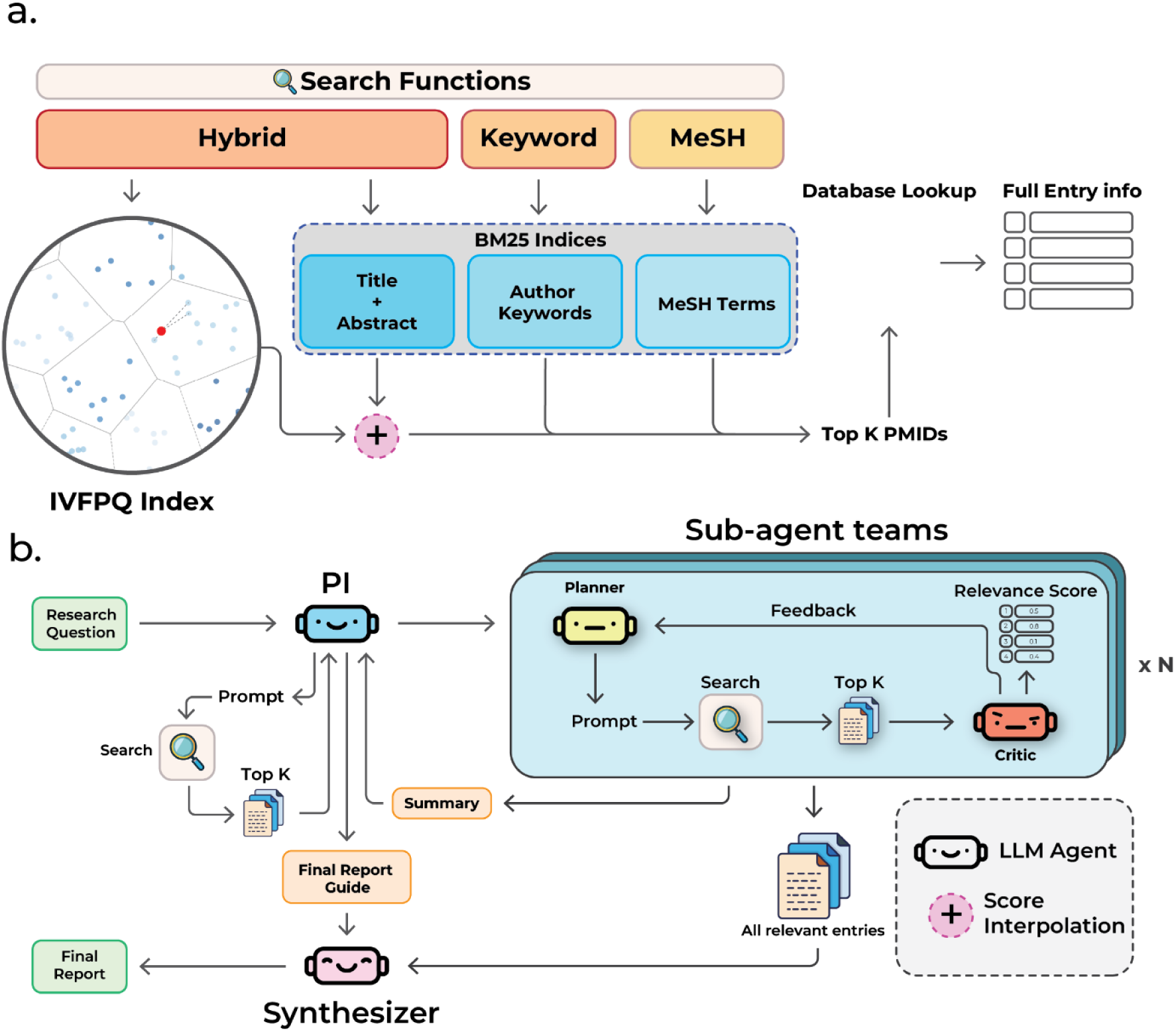
Retrieval engine and agentic orchestration. **a.** Hybrid retrieval over the entire PubMed-derived corpus (28.3M title–abstract passages). A FAISS IVFPQ index provides dense semantic search, while three BM25 indices cover title+abstract, author keywords, and MeSH. Candidates are merged via score-level interpolation to produce top-K PMIDs, followed by database lookup for full metadata. **b**. Multi-agent workflow. A Principal Investigator (PI) delegates to planner–critic sub-teams that iteratively search, score relevance, and refine evidence; a synthesizer compiles the final, citation-grounded report. The system adapts search functions (Hybrid, Keyword, MeSH) per task and logs all steps for reproducibility.

We selected Inverted File with Product Quantization (IVFPQ) [23] as our indexing strategy. IVFPQ operates in two stages: first, an inverted file structure partitions the vector space into coarse clusters, then product quantization compresses vectors within each cluster. During index construction, we trained a quantizer to divide the 4,096-dimensional embedding space into 16,384 coarse clusters using k-means clustering on a representative training sample. This training sample was collected by randomly sampling 1 million of the 28.3 million embeddings, yielding sufficient coverage of the vector space while remaining within our memory budget. After training the coarse quantizer, we applied product quantization within each cluster, decomposing the 4,096-dimensional space into 64 subspaces of 64 dimensions each, then quantizing each subspace independently with 8-bit codes. This compression reduces memory footprint from 8 kilobytes per vector (4096 dimensions × 2 bytes) to approximately 64 bytes (64 subspaces × 1 byte), enabling the entire index to fit in memory for fast query processing. At query time, we set nprobe=492 (approximately 3% of clusters), directing the search to examine 492 of the 16,384 clusters. This configuration balances retrieval quality with query latency, enabling sub-second search across the entire corpus.

In parallel with the FAISS index, we constructed three Best Match 25 (BM25) indices for lexical retrieval [24]. BM25 is a probabilistic ranking function that scores documents based on term frequency and inverse document frequency, with saturation to prevent over-weighting of repeated terms. Unlike dense retrieval, which captures semantic similarity through learned embeddings, BM25 provides exact term matching with interpretable relevance scores, making it particularly effective for queries with specific technical terminology or rare terms that may not be well-represented in the embedding space.

We built three separate BM25 indices using the BM25S library [25] with over different textual fields. The first index covers concatenated title and abstract text, combining both fields with a space separator to provide broad semantic matching through term statistics. The second index targets author-assigned keywords, which often capture domain-specific terminology more precisely than free text. The third index operates over Medical Subject Headings (MeSH) terms, the controlled vocabulary used to index biomedical literature in PubMed [1]. This multi-view lexical retrieval exploits the structured organization of biomedical literature, where curated vocabularies frequently provide more precise concept matching than uncontrolled natural language.

Agents interface with three retrieval functions: Hybrid Search, Author Keyword search and MeSH term search (**Fig. 1b**). The hybrid search function executes parallel retrieval against the FAISS vector index and a BM25 index over title and abstract fields. Initially, we retrieve top k*100 candidate papers, where k is the number of papers and then merge them using score-level interpolation, where normalized FAISS similarity scores and BM25 relevance scores are combined via a tunable weighting factor α. This weighting allows the model to control the contribution of semantic similarity versus keyword overlap depending on the query type or task objective.

In addition to the hybrid search, dedicated MeSH term and author keyword search functions operate over their respective BM25 indices, enabling agents to perform vocabulary-controlled retrieval when standardized terminology or domain-specific keywords provide better precision.

### Agent Orchestration

The system is organized as a hierarchical multi-agent architecture designed to decompose complex scientific questions into smaller, verifiable reasoning tasks (**Fig. 1b**). This design is grounded in a consistent observation about large language models: while they perform individual, well-scoped reasoning steps reliably, their accuracy and interpretability decline when confronted with long, unstructured contexts or multi-objective tasks [26, 27]. Queryome addresses this by converting literature analysis into a distributed reasoning process, where multiple specialized agents collaborate through well-defined roles of investigation, critique, and synthesis.

At a conceptual level, a central coordinating Principal Investigator (PI) agent (blue) interprets the research question, formulates an initial search strategy, and decides whether the problem should be divided into multiple investigative threads. Each thread is executed by a planner (yellow)–critic (red) pair operating in iterative cycles. The planner designs retrieval queries and calls external search tools, while the critic evaluates the relevance and quality of the retrieved material, assigns scores with justification, and guides further refinement. This feedback loop transforms retrieval into a reasoned exploration rather than a one-shot lookup. Once sufficient evidence has accumulated, each team produces a concise summary, which is later integrated into a broader synthesis.

The workflow itself is agent-controlled, enabling Queryome to dynamically scale the orchestration depth and breadth according to the complexity of the query (**Fig 2**). Simple factual questions may be resolved by a single agent executing a limited retrieval cycle, whereas open-ended biomedical inquiries can trigger the autonomous creation of multiple planner–critic teams operating in parallel. This design allows the system to allocate reasoning effort proportionally to problem difficulty, expanding when interpretive depth is needed, contracting when a direct answer suffices.

**Figure 2.**
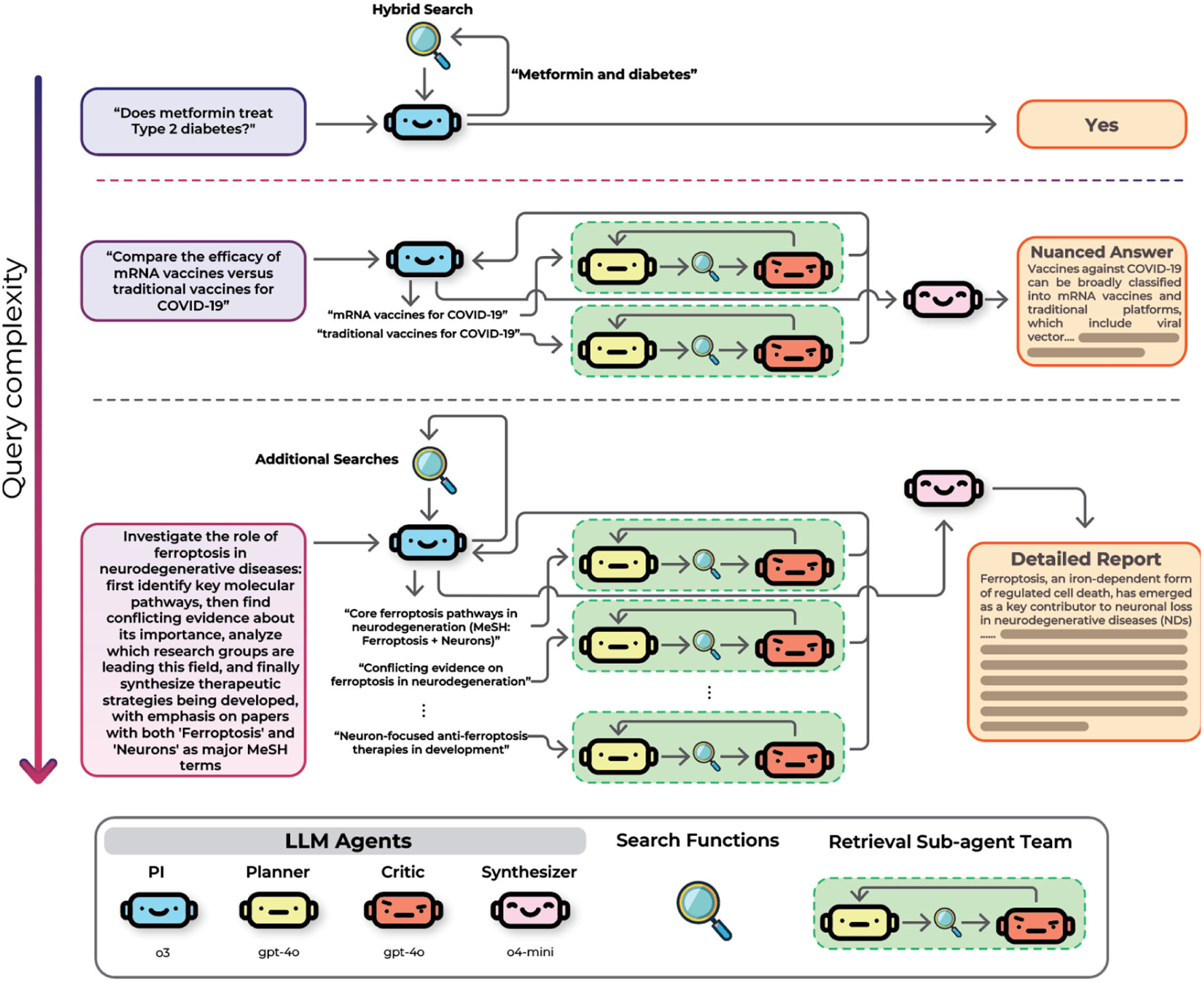
Scaling behavior with query complexity. The PI routes simple fact-checks to a single hybrid search, yielding a direct answer. Comparative questions trigger multiple planner–critic loops that search in parallel and return a nuanced synthesis. Open-ended research prompts (e.g., pathway mapping, controversy analysis, labs/therapies landscape) spawn additional searches and multiple sub-teams; their summaries are consolidated into a detailed, evidence-grounded report. Agent roles: PI (GPT-o3) for reasoning, planner and critic (GPT-4o) for retrieval and evaluation, and synthesizer (GPT-o4-mini) for final assembly. Top, in this simple query of a drug for type 2 diabetes, the PI performs a single hybrid search and provides a direct answer. Middle, in the question that asked to compare between vaccine types for COVID-19, two subagents investigated the evidence for mRNA and traditional vaccines, respectively. The results were then combined into a more nuanced answer by the synthesizer. Bottom, in this open-ended research question on ferroptosis, multiple subagent teams were spawned to investigate different sub questions with different search strategies as instructed by the initial prompt. The summaries and retrieved papers were then compiled into a review style response by the synthesizer.

In our implementation, the PI uses o3 for multi-step inference and tool-based reasoning. Planner and critic agents use GPT-4o, and the synthesizer employs o4-mini. We selected o3 for the PI because our experiments demonstrate that reasoning models extract far more value from the multi-agent architecture than standard models. GPT-4o is used for planner and critic agents because it supports native tool calling. Native tool calling of LLMs is an ability to access external information by generating schema constrained text that can be passed as arguments to arbitrary function or code. The GPT-4o model supports this ability natively, i.e., no additional finetuning is necessary. In Queryome, tool calling has been extensively used in database search, based on the question context, the LLM generated arguments for procedural calls to retrieve information from the RAG database. For the synthesizer, o4-mini is fast, inexpensive, and supports long context lengths, suiting its role of compiling retrieved evidence into a final response. All prompts, retrieval calls, critic evaluations, and outputs are logged for reproducibility; full system prompts are provided in Supplementary Information 1. This modular, self-scaling design embodies the guiding philosophy behind Queryome: delegate narrowly, reason explicitly, and synthesize only from verified evidence. By transforming biomedical research into a network of cooperative, high-confidence reasoning steps, Queryome achieves both scalability and interpretability that traditional single-pass LLM pipelines cannot sustain.

## Results

We evaluated Queryome across multiple dimensions to comprehensively assess its capabilities. First, we tested Queryome on MIRAGE (Medical Information Retrieval-Augmented Generation Evaluation), a large-scale biomedical question-answering benchmark comprising 7,663 questions from five established datasets [13]. We also examined Queryome’s instruction-following ability through its performance on PubMedQA [28], where we evaluated whether the system could adapt its evidence synthesis strategy based on task-specific requirements. To assess genuine reasoning capabilities beyond pattern matching, we evaluated Queryome on the biomedical subset of the Humanity’s Last Exam (HLE) benchmark [19], which requires multi-step compositional reasoning over long-form questions. Finally, we compared Queryome’s knowledge extraction and synthesis capabilities against state-of-the-art commercial deep research tools through a Review Generation Test that measured the system’s ability to reconstruct expert-level scientific reviews.

### Benchmark results on MIRAGE

We evaluated Queryome on the MIRAGE benchmark, which was specifically created to systematically evaluate RAG systems for medical question answering, addressing the challenges of hallucinations and outdated knowledge that plague standalone large language models [13]. The benchmark’s diversity across medical examination formats and biomedical research questions makes it a comprehensive test of both retrieval effectiveness and reasoning capabilities across different types of medical information needs.

MIRAGE comprises 7,663 questions from five commonly used biomedical QA datasets [12]: Massive Multitask Language Understanding Medical (MMLU-Med) subset [29] with 1,089 questions covering anatomy, clinical knowledge, professional medicine, human genetics, college medicine, and college biology, MedQA-US [30] with 1,273 four-option questions from the US Medical Licensing Examination, MedMCQA [31], which is medical multiple-choice questions from Indian medical entrance exams, PubMedQA [28], which is biomedical research questions with yes/no/maybe answers, and BioASQ-Y/N [32], which includes 618 yes/no questions from biomedical research spanning 2019-2023. Thus, the benchmark includes questions ranging from medical examination formats that test clinical knowledge and reasoning to biomedical research questions that require synthesis of scientific literature.

Table 1 shows the summary of the benchmark. We allowed Queryome to deploy up to 10 parallel subagent teams, with each team conducting iterative planner-critic cycles to gather evidence from our PubMed-based retrieval system. Each subagent team was allowed to perform a maximum of 5 retrieval iterations or until the critic agent determined sufficient evidence had been gathered. We ran Queryome three times and reported standard deviation to capture variability from model. This randomness comes from sampling the predicted token probability distribution during generation. Most models allow temperature control where for evaluations this parameter is often set to zero to produce a deterministic output, but OpenAI’s o3 does not expose a temperature setting, so repeated runs are conducted to quantify its variance.

**Table 1.** Summary of the Benchmark on MIRAGE.

| Method | PubMedQA | BioASQ | MMLU | MedQA | MedMCQA | Average |
| --- | --- | --- | --- | --- | --- | --- |
| GPT-4 + MedRAG | 70.6 | **92.56** | 87.24 | 82.8 | 66.65 | 79.97 |
| Llama-3 + MedRAG | 71.6 | 89.97 | 85.58 | 76.9 | 68.87 | 78.59 |
| Llama-3 + CoT | 59 | 83.01 | 85.77 | 80.91 | 70.93 | 75.92 |
| SimRAG 8B | 80 | 91.75 | 75.57 | 62.92 | 67.51 | 75.55 |
| SimRAG 27B | 73.6 | 92.07 | 81.63 | 63.63 | 64.16 | 75.02 |
| Queryome | **81.67 ± 0.23** | 92.39 ± 0.24 | **95.84 ± 0.31** | **93.50 ± 0.48** | **81.48 ± 0.23** | **88.98 ± 0.14** |

The percentage of questions the methods answered correctly is shown. The highest value observed for each category is shown in bold. We ran Queryome three times and report the average and the standard deviation from the runs. The results of the five other systems are taken from the MIRAGE paper. These are the top performing systems.

**Table 2.** Performance summary on the biomedical subset of Humanity’s Last Exam.

| Model | Accuracy (%) | Calibration Error (%) |
| --- | --- | --- |
| Queryome | $24.9 \pm 0.6$ | $59.2 \pm 1.9$ |
| GPT-5 | $19.8 \pm 0.9$ | $59.9 \pm 0.8$ |
| O3 | $14.6 \pm 2.3$ | $53.8 \pm 3.6$ |

Accuracy is the proportion of questions answered correctly, and Calibration Error is the root-mean-square (RMS) calibration error between predicted confidence and outcome. All systems were evaluated with three replicates. Detailed performance metrics for each replicate are provided in Supplementary Tables 1-3.

We compared Queryome with five other approaches reported as state-of-the-art methods in the MIRAGE paper: GPT-4 + MedRAG, Llama-3 + MedRAG, Llama-3 + CoT, SimRAG-8B, and SimRAG-27B [13]. The prompts used for each MIRAGE dataset are provided (Supplementary Information 2). As shown in Table 1, Queryome showed overall the highest average accuracy of 88.98% with a large margin to the second method, GPT-4 + MedRAG (79.97%). Among the five datasets, Queryome had the highest accuracy for four datasets except for BioASQ. The performance gains by Queryome were particularly pronounced on reasoning-intensive medical examination tasks. On MMLU-Med, Queryome scored 95.84% ± 0.31%, and on MedQA-US, we achieved 93.50% ± 0.48%. While these substantial improvements on medical licensing examination questions may be attributable to the enhanced reasoning capabilities of the o3 model underlying our Primary Investigator agent, the gains extend beyond pure reasoning ability as we discuss later in the Ablation studies section. Queryome’s architecture enables it to retrieve and synthesize evidence from multiple sources to assess scientific consensus, rather than relying on parametric knowledge alone. This strength is evident across the biomedical research tasks as well: on BioASQ-Y/N, Queryome scored 92.39% ± 0.24%, on PubMedQA we attained 81.67% ± 0.23% using our modified instruction-following approach, and on MedMCQA, Queryome reached 81.48% ± 0.23%. The system’s ability to identify converging evidence across the literature, reconcile contradictory findings, and ground answers in peer-reviewed sources distinguishes it from traditional biomedical RAG systems.

### Remarks on PubMedQA and Instruction Following

The agentic paradigm of Queryome facilitates the exploration and integration of diverse, complementary, and even contradictory concepts across the RAG database. This approach enables the system to address more complex problems and expands the reasoning-based knowledge capabilities of Queryome. Interestingly, we noticed that this broader scope led to a divergence from the ground-truth labels provided in PubmedQA. With the prompts we originally used (Supplementary Information 1.2.1), the accuracy recorded for Queryome achieved was 72.4%. While examining “failed” cases, we found that a different answer from the “ground-truth” was yielded by the Queryome system’s adherence to the broader scientific consensus retrieved from multiple papers, which seemed to be more correct, compared to the “ground-truth” answers generated from a rather simplistic protocol used in PubmedQA to generate an answer from the abstract of a single paper.

For example, for a question that asked whether oral endotracheal intubation is effective in helicopter environments, Queryome cited a 2024 systematic review [33] and large observational studies such as Harrison et al. 1997 [34] showing equivalent success rates between helicopter and ground settings. Based on this evidence, the Queryome’s final response was “no,” which is well justified by the current literature. However, the PubMedQA benchmark labeled the correct answer as “yes,” because it relies on an older manikin simulation study by Stone and Thomas 1994 [35] whose findings are contradicted by later field data. Similar discrepancies appeared in other cases. For instance, when asked about the Barthel Index as a measure of long-term stroke outcomes, Queryome cited longitudinal evidence of persistent cognitive and psychosocial deficits (Wilkinson et al., 1997; Lai et al., 2002; van Exel et al., 2004) [36–38] and answered “no.” Yet PubMedQA marked “yes” as correct, based on the single earlier study by Wilkinson et al. 1997 [36].

The same types of mismatches between evidence-based reasoning and the benchmark’s ground-truth labels were frequent. Further investigation of the PubMedQA benchmark clarified the source of this discrepancy. The dataset was constructed by pairing each question with a PubMed article title, providing the abstract shorn of its concluding sentence as context, using the concluding sentence itself as the long-form answer, and distilling that conclusion into a yes/no/maybe label [28]. As a result, the benchmark is primarily oriented toward information retrieval and extraction rather than reasoning.

To reflect the way questions and answers are crafted in PubMedQA, we ran the experiment with a modified instruction prompt (Supplementary Information 2). This information of how the benchmark is constructed is available in the PubMedQA paper [28]. Other conditions were unchanged. With this adjustment, Queryome aligned more closely with the benchmark’s intent. With this prompt, the accuracy of Queryome improved to 81.67 % ± 0.23 %, surpassing the reported state-of-the-art. This result demonstrates not only Queryome’s information retrieval and extraction capabilities but also its strong instruction-following ability.

### Benchmark results on Humanity’s Last Exam

To test whether Queryome’s performance gains reflect genuine evidence-based reasoning rather than an easy benchmark or model overfitting, next we evaluated the system on a more demanding test. We turned to HLE, a challenging benchmark developed by the Center for AI Safety and Scale AI consisting of approximately 3,000 expert-level questions across mathematics, humanities, and natural sciences [19]. HLE was specifically designed to address benchmark saturation, where state-of-the-art models now achieve over 90% accuracy on popular benchmarks like MMLU [29], and features questions crowdsourced from nearly 1,000 subject matter experts.

We restricted our evaluation to HLE’s biomedical, text-only subset, which consists of questions spanning molecular biology, physiology, biochemistry, pathology, and clinically oriented biomedical reasoning. The prompts used for both base LLM evaluation and Queryome on HLE are provided in Supplementary Information 3. This resulted in a benchmark consisting of 222 questions. We test Queryome, relative to o3 and GPT-5 on this subset on September 22, 2025. We compared with o3 as it is the base model used by the PI in Queryome. And GPT-5 was the top model on the Human’s Last Exam at the time of our development. In addition to reporting accuracy on the public subset of HLE questions, we utilized the Root Mean Square (RMS) calibration error to evaluate the reliability of model confidence [39]. Conceptually, a perfectly calibrated model should exhibit confidence levels that mirror its actual accuracy; for instance, predictions made with 50% confidence should be correct 50% of the time. Significant deviations from this ideal. particularly instances of high confidence paired with low accuracy, serve as quantitative indicators of confabulation. In our initial analysis, we observed that most models displayed systematic calibration errors exceeding 0.50 alongside accuracies below 10%, suggesting a pervasive tendency toward hallucination on these expert-level tasks.

To contextualize Queryome’s performance, we constructed baselines using state-of-the-art reasoning models evaluated without access to our specialized paper retrieval pipeline. Among the three methods, Queryome achieved the highest average score of 24.9%. The base GPT-5 model showed an accuracy of 19.8 ± 0.9% with a calibration error of 59.9%. The o3 model, which serves as the reasoning core for Queryome’s PI agent, had a baseline accuracy of 14.6 ± 2.3% with a calibration error of 53.8%. While Queryome’s calibration error (59.2%) remains comparable to the base models reflecting the broader challenge of overconfidence in LLMs on expert tasks, the substantial increase in accuracy demonstrates the efficacy of grounded retrieval in resolving complex scientific queries.

To better understand the source of these improvements, we analyzed specific instances where Queryome’s evidence-driven reasoning enabled correct answers that the base reasoning model missed. For example, Question 13 asks: "Which animal might be able to hear human muscle twitches?" The base reasoning model incorrectly answered "Bat," assuming that bats’ ultrasonic hearing (up to 100 kHz) would allow them to detect muscle activity. However, Queryome correctly identified the answer as "Whale." Through six iterative searches, the system retrieved evidence establishing that human skeletal muscle acoustic energy is concentrated below ∼20 Hz (PMID 2710145) and that baleen whales possess infrasonic hearing capabilities down to ∼10 Hz (PMID 17516435). By integrating human biomechanics literature with comparative auditory physiology, Queryome successfully corrected the model’s erroneous assumption about frequency ranges.

Similarly, Question 92 asks: "What lab parameter could best indicate the cause of rapid renal function decline in a patient with SLE who developed throat infection after steroid discontinuation?" The base model incorrectly selected "rising anti-dsDNA antibody titer," anchoring on the SLE diagnosis to infer lupus nephritis activity. While logical on the surface, this failed to account for the specific clinical presentation. Queryome correctly identified "Antistreptolysin-O (ASO) titers" as the answer. By retrieving literature on post-infectious glomerulonephritis in immunocompromised adults (PMID 2006561, PMID 23691638), the system recognized that the three-week latency between the throat infection and renal decline was characteristic of a superimposed streptococcal infection rather than a lupus flare, demonstrating the value of precise, context-aware retrieval.

### Performance on a Review Generation Test

Next, we sought to assess how the system performs relative to existing commercial deep research tools in real-world scientific synthesis tasks. To address this, we compared Queryome against four state-of-the-art AI research systems including OpenAI Deep Research [16], Gemini Deep Research [18], Perplexity Deep Research, and Scite.AI [40].

We developed a review generation test to evaluate each system’s ability to synthesize comprehensive scientific reviews (**Fig. 3a**), which is formulated as follows: First, we selected five recently published review articles from diverse biomedical domains that are widely regarded as high-quality syntheses of their respective fields. Specifically, we selected: a review on integrating spatial transcriptomics with tissue morphology, which proposes a translation-integration framework for combining imaging-derived features with spatial gene-expression data [41]; a review on biomonitoring and wearable technologies for women’s health, detailing emerging sensors for fertility tracking, pregnancy monitoring, cancer detection, and hormone measurement [42]; a review on translational regulation during cellular stress, summarizing how proteotoxic, oxidative, and nutrient-sensing stresses reshape global and selective protein-synthesis programs and influence ageing [43]; a review on type-2 immunity in cancer, reassessing TH2- and ILC2-driven pathways and describing conditions where type-2 responses may suppress or constrain tumor growth [44]; and a review on long-term evolutionary studies, highlighting how decades-scale field and laboratory systems reveal evolutionary dynamics, innovation events, and speciation processes invisible to short-term studies [45]. For each review, we analyzed the paper to extract its core research question and scope, then formulated a standardized prompt instructing AI systems to generate a comparable review de novo. Each deep research system received identical prompts and was allowed to use its native retrieval and synthesis pipeline without constraints. The prompts for each review are provided in Supplementary Information 4.

**Figure 3.**
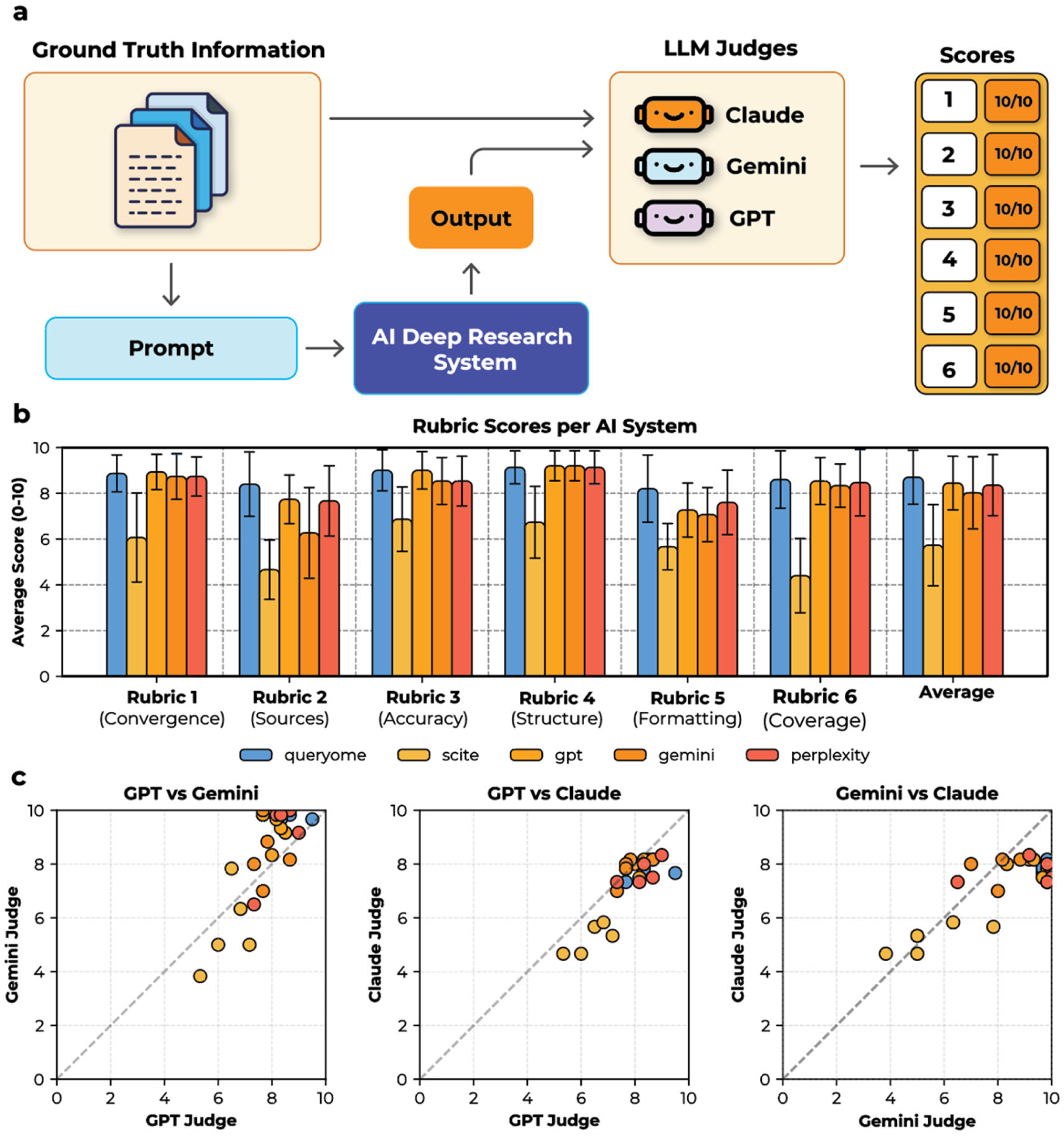
Results of the review generation test. **a.** Evaluation setup: each system generates a literature review for a target topic; three LLM judges (Claude, Gemini, GPT) independently score six rubrics (content convergence, source quality & coverage, factual accuracy, structure, formatting, and comprehensiveness). **b.** Rubric scores with means and standard deviations across the three judges. Five systems were compared, Queryome (blue), Scite.AI (yellow), GPT Deep Research (orange), Gemini Deep Research (dark orange), and Perplexity Deep Research (red). **c.** Pairwise comparisons of average scores by the three judges. The color of data points indicates the system that was evaluated. The color code is the same as the panel b. The correlation between GPT and Gemini is 0.69; GPT vs Claude: 0.65; Claude vs Gemini: 0.61

For evaluating free text of the generated review, traditional text similarity metrics such as ROUGE [46] and BLEU [47] are inadequate, as they reward surface-level lexical overlap rather than conceptual accuracy, reasoning depth, or synthesis quality. We therefore opted for LLM-based evaluation, where language models are used to compare AI-generated reviews against the original published reviews as the ground truth and score them. To minimize bias and subjectivity, we employed three independent LLM judges: Claude 4.5 Sonnet (Anthropic) [48], Gemini 2.5 Pro (Google DeepMind) [49], and GPT-5 (OpenAI) [50]. We prompted each judge to evaluate the following six rubrics, with each scored from 1 (worst) to 10 (best):

Content Convergence, which quantified the alignment between the generated review and the reference article in terms of covered topics, central findings, and articulated conclusions, including identification of the same domain-specific trends, gaps, and insights. Source Quality and Coverage assessed whether the model cited credible scholarly literature, recovered key references appearing in the ground-truth review, and incorporated additional high-value sources where appropriate. Factual Accuracy measured the correctness of all scientific statements and verified the absence of hallucinated mechanisms, misinterpretations, or unsupported claims. Structure and Organization evaluated the logical sequencing of ideas, coherence of the narrative, and appropriateness of section boundaries. Formatting and Presentation measured adherence to academic writing conventions, clarity of exposition, and consistency in citation style. Finally, Comprehensiveness quantified the breadth and depth of topic coverage, including whether the model addressed all major subtopics and relevant conceptual perspectives present in the reference. The prompt provided to the judges is shown in Supplementary Information 5. Each judge provided scores with brief justifications for each rubric. We report mean scores across the three judges with standard deviations to capture inter-judge variability.

**Fig. 3b** summarizes the scores of the five systems including Queryome. Queryome achieved the highest average score of the six rubrics (52 ± 1.6, the average and the range of the scores by the three judges), indicating superior faithfulness to the original review. Among the baseline systems, OpenAI Deep Research and Perplexity Deep Research came the second (≈ 50 ± 1–4), and Gemini Deep Research followed closely (48.1 ± 2.9). SciteAI, which relies primarily on citation graph reasoning, lagged significantly (34.4 ± 5.0) in reconstructing conceptual linkages and thematic flow. As shown in **Fig. 3c**, the evaluations by the three judges were fairly consistent. The correlation coefficients between pairs of judges ranged from 0.61 to 0.69. Correlations of judges’ evaluations per rubric is provided in Supplementary Figure 1. Complete scoring data including mean and standard deviation per rubric are provided in Supplementary Tables 4–6, and raw scores by each judge for all five target papers are provided in Supplementary Table 7. Links to the original evaluation conversations are available in Supplementary Table 9.

Queryome showed the highest score in Rubric 2 (Source Quality & Coverage) and Rubric 5 (Formatting & Presentation) with relatively large margins. The reason behind these results is perhaps the system’s agentic architecture. Unlike standard web search which often relies on keyword matching, Queryome’s hybrid retrieval enables the discovery of conceptually related but lexically distinct evidence. Furthermore, the iterative planner-critic loops may be able to facilitate an exploratory search strategy where every retrieved abstract is explicitly reasoned about before being accepted as evidence.

Queryome demonstrated strong alignment with the ground truth by accurately capturing key mechanistic insights. For example, Paper 4 described how tumor-specific IgE binding to FcεRI+ mast cells can enhance dendritic cell antigen uptake and promote CD8+ T cell activation; a mechanism central to the emerging field of allergooncology [44]. Queryome found information regarding this pathway and noted that "when ligated by tumor-specific IgE, FcεRI+ mast cells release TNF-α and leukotrienes that enhance antigen uptake by dendritic cells and promote CD8 activation." In contrast, competing systems discussed mast cells primarily in pro-tumorigenic terms or omitted this IgE-mediated crosstalk entirely, missing a key therapeutic avenue emphasized in the ground truth.

As an agentic system with semantic search capabilities, we expect Queryome to find gaps in evidence and make connections to literature outside the scope of the original review. We examined Queryome’s reproduction of Paper 2 and found notable examples of such behavior. Paper 2 discussed emerging wearable biomonitoring technologies that aim to improve diagnosis, tracking and equitable access to care for women’s health conditions [42]. In the original review (Paper 2), the authors briefly noted privacy concerns regarding reproductive data, recommending "on-device encryption" as a potential safeguard. However, the text treated this solution as a theoretical ideal rather than a technically validated approach. Queryome’s planner-critic architecture identified this lack of concrete evidence as a gap. To address it, the system retrieved and synthesized two distinct types of sources. First, it identified Alfawzan & Christen’s (2023) comprehensive analysis of FemTech ethics [51], providing the necessary regulatory context. Second, and crucially, it discovered Tjhin et al.’s (2025) implementation study on user-friendly differential privacy applications [52]. By integrating these sources, Queryome transformed a high-level recommendation into a concrete engineering pathway, effectively bridging the gap between abstract problem identification and technical feasibility. In contrast, privacy concerns were acknowledged but not operationalized in the responses from the other systems. OpenAI and Gemini reiterated high-level risks without specifying technical mechanisms; Perplexity did not retrieve relevant differential-privacy implementations; and Scite.AI offered only generalized statements on “secure data.”

Furthermore, the system demonstrated cross-domain reasoning regarding access equity, highlighting the specific utility combining an agentic architecture with semantic search. While the ground-truth review acknowledged that "broadband access" limits wearable adoption in low-income settings, it stopped short of offering solutions or in-depth analysis. Queryome enriched this narrative by identifying and citing Ebekozien et al.’s (2024) study on health inequities in diabetes care [53]. This retrieval is significant because a standard keyword search for "fertility wearables" likely would not have surfaced a paper on diabetes. However, Queryome’s semantic search recognized that the infrastructure barriers described in the diabetes literature such as the need for consistent connectivity and device literacy, are conceptually identical to those facing women’s health technologies. This allowed the system to perform sophisticated cross-domain reasoning, transferring insights from a more mature digital health field to address systemic "digital divide" challenges in a new context which the other Deep Research systems did not mention.

### Ablation Studies

We conducted ablations on MIRAGE to evaluate how model choice affects performance (Fig. 4). These experiments test two hypotheses: first, that strengths of the language model, namely, native tool-calling capability, matters more than model scale for agentic architectures; and second, that reasoning models are necessary to effectively coordinate multi-step evidence gathering.

**Figure 4.**
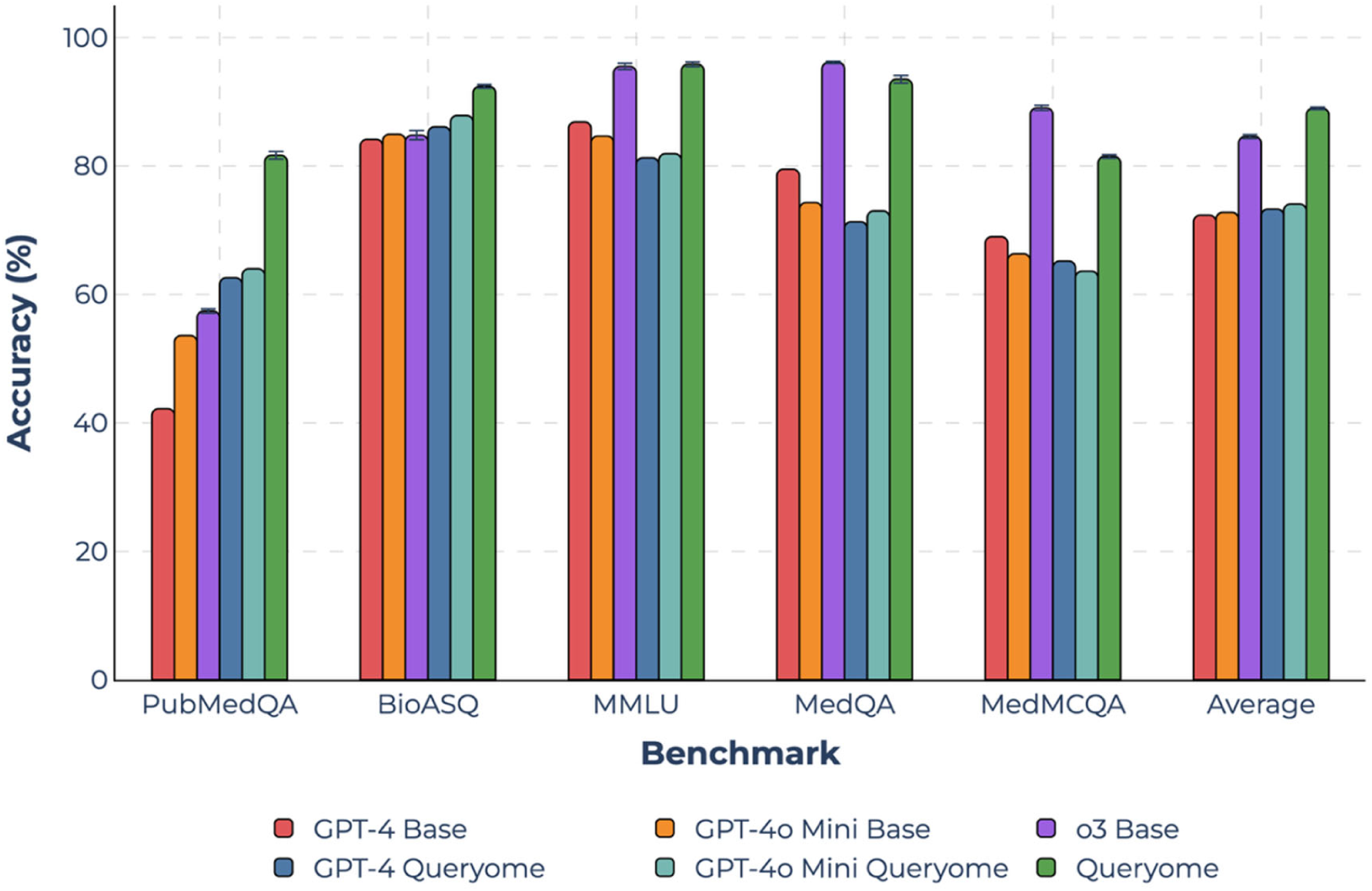
Results of ablation study on MIRAGE. We compared Queryome with five other conditions: three base models (GPT-4, GPT-4o-mini, and o3) and two alternative Queryome configurations where all agents use the specified model family, GPT-4 or GPT-4o. Accuracy is the percentage of the correctly answered questions. For GPT-4 and GPT-4o-mini conditions, we report single runs with temperature set to 0 for deterministic output. For o3 conditions, the mean of three replicates are reported because the temperature parameter is not exposed to the user. The error bars indicate standard deviation. Average accuracy across all MIRAGE benchmarks: GPT-4 Base: 72.3%; GPT-4o-mini Base: 72.8%; o3 Base: 84.6%; GPT-4 Queryome: 73.3%; GPT-4o-mini Queryome: 74.1%; Queryome (o3 as PI): 89.0%. Links to benchmark responses are provided in Supplementary Table 10.

We evaluated six configurations. "Base" refers to the LLM answering questions as a standalone model, i.e., they were used directly without retrieval or multi-agent orchestration. We chose three “base” models of differing nature, scale and capability for comparison [67]: GPT-4, a strong general-purpose model; GPT-4o-mini, which is a lightweight model optimized for efficiency; and o3, a model specialized for deep thinking and reasoning. As mentioned earlier, Queryome uses o3 for the PI agent.

GPT-4 and GPT-4o-mini are standard instruction-following models, whereas o3 is a reasoning-focused model. Standard models can perform multi-step tasks but are not explicitly trained to improve chain-of-thought reasoning. On the contrary, reasoning models are post-trained via reinforcement learning to enhance deliberative, multi-step thinking and reasoning. Moreover, GPT-4o-mini and o3 support native tool calling capabilities while GPT-4 does not.

Additionally, we included Queryome and its two variants. For the standard Queryome configuration, o3 serves as PI, GPT-4o handles the planner and critic roles, and o4-mini serves as synthesizer. For the ablated variants, GPT-4 Queryome and GPT-4o-mini Queryome, all agents use the specified model. Thus, in GPT-4 Queryome, the planner and critics are on par, considering GPT-4 and GPT-4o are almost the same models, and synthesizer is stronger than in the standard Queryome. In the GPT-4o-mini Queryome, the synthesizer is the same and planners and critics are weaker than the standard Queryome. Given that the sub-agents are assigned the simpler roles of retrieval and synthesis, utilizing the same model as the PI is unlikely to affect the overall orchestration.

First, we compare base models. In all five datasets, o3 showed the highest accuracy (84.6% average vs 72.3% for GPT-4 and 72.8% for GPT-4o-mini). This demonstrates the effectiveness of explicit post training for reasoning as we see that o3 particularly dominates exam-style benchmarks (MMLU, MedMCQA, MedQA). Interestingly, GPT-4o-mini outperformed GPT-4 on the two evidence-grounded tasks (PubMedQA: 53.6% vs 42.2%; BioASQ: 85.0% vs 84.1%). This result is counterintuitive, as model scale is generally expected to dominate the memorization capabilities being assessed in these benchmarks. However, because these models are effectively "black boxes" with proprietary training strategies and datasets, determining the precise cause of this disparity is non-trivial. Potential factors may include but not limited to differences in data quality and distribution, the extent of instruction tuning, or even the internal use of tool-calling mechanisms.

Next, we compare the Queryome variants. Overall, the standard Queryome performs the best (89.0% average), and GPT-4o-mini Queryome outperforms GPT-4 Queryome on four out of five benchmarks. The comparison between GPT-4o-mini and GPT-4 in the Queryome setting is noteworthy because GPT-4o-mini is capable of both performing evidence retrieval through tool-usage and answering in a structured format in an agentic setting without manual orchestration while having less parameters. The advantage of the standard Queryome over GPT-4/GPT-4o-mini Queryome is considerable on PubMedQA, MedQA, and MedMCQA, but less pronounced on BioASQ. BioASQ is a knowledge-based, binary yes/no benchmark with already a high base accuracy (84–85%), leaving less headroom of improvement. Critically, from our analysis, we observe that BioASQ questions are often resolved within shallow evidence gathering (generally one retrieval call per question), so there is less need for PI-level multi-step planning and reasoning, as performance is driven by the breadth and quality of the knowledge base, rather the competency of the model.

Finally, we address the critical question of whether the improved performance is attributable to the base o3 reasoning model or to Queryome’s novel agentic paradigm. On average, Queryome outperforms the o3 base model on average (89% vs. 84.6%). Specifically, Queryome excels in PubMedQA (+24.3 percentage points, pp) and BioASQ (+7.6 pp), as these benchmarks are directly knowledge-based. Consequently, Queryome’s access to a well-curated biological literature database and its ability to retrieve information proves far more effective than relying solely on internal memorization, as is the case with o3. Conversely, for the remaining benchmarks, the performance gains were marginal (+0.3 pp on MMLU) or even negative (−2.6 pp on MedQA and −7.6 pp on MedMCQA). An analysis of the reasoning traces in cases where o3 was correct but Queryome was wrong reveals two primary factors. Firstly, Queryome is optimized and explicitly instructed to follow retrieved evidence. However, both datasets often reward "curriculum heuristics" and "single-best-step" conventions that can diverge from nuanced information found in the literature. For instance, Queryome favored evidence-supported scabies mass drug administration over the MedMCQA key (Q1535).

Similarly, in MedQA Question (Q402), a conflict arises between empirical evidence and clinical exam logic regarding emergency contraception. It is a multiple-choice question, asking suitable emergency contraception options for a young woman with metal allergy, and the choices are copper intrauterine device (Cu-IUD), mifepristone, ulipristal acetate, and depot medroxyprogesterone acetate. Through retrieval, Queryome identified that copper allergies are clinically rare (Goldstuck & Cheung, 2019 [68]), leading to the conclusion that a metal allergy should not necessarily contraindicate the use of Cu-IUD, thus chose Cu-IUD as the answer. However, the designated correct answer is “administering ulipristal acetate”, prioritizing clinical risk mitigation. In a standardized exam context, a documented allergy acts as a specific constraint, favoring the safer pharmacological alternative, ulipristal acetate. While the o3 model correctly identified Ulipristal by strictly adhering to these exam heuristics, Queryome’s reasoning demonstrates a high level of contextual awareness. Thus, Queryome’s answer would be also logical, because it validated the safety of the most effective treatment (Cu-IUD) before recommending it, prioritizing clinical efficacy over the likely nominal risk of the stated allergy.

Secondly, even when retrieved snippets are relevant, errors are occasionally amplified by noise, i.e., ambiguous, conflicting or spurious retrievals, in Queryome, particularly when distractors in the question are partially supported by the text. While it is debatable whether the o3 model is optimized for MedQA and MedMCQA, given their popularity in LLM ranking, empirically it turns out that these noises influenced the inherent memorization of our PI model, as without the literature grounding the PI model, i.e., the vanilla o3 model could answer them correctly. The limitation of retrieving only from abstracts rather than full-text articles contributes to this noise. Access to the full texts would likely enable more grounded knowledge discovery and deep reasoning. Regardless, the HLE benchmark provides the most rigorous assessment of competency, as it rewards multi-stage reasoning over simple knowledge retrieval or memorization. On this benchmark, Queryome demonstrated a substantial relative accuracy improvement of over 70% compared to o3. In summary, although Queryome utilizes o3 as its primary model, our agentic orchestration considerably enhances the base model’s capabilities through the simultaneous aggregation of retrieval, reasoning, and evidence-guided decision-making.

## Discussion and Conclusion

Biomedical literature continues to expand at a rate that outpaces human capacity for synthesis, demanding AI systems that can reason scientifically rather than merely retrieve. The results of this work show that meaningful progress in biomedical question answering depends not on retrieval scale but on the integration of structured reasoning with domain-specific evidence. Queryome demonstrates that combining multi-agent coordination, explicit evidence evaluation, and specialized retrieval produces measurable improvements in factual grounding, synthesis coherence, and reasoning depth across diverse biomedical tasks.

The improvement arises when retrieved evidence is interpreted through reasoning, not when it is simply appended to a prompt. In tasks demanding causal inference, temporal reasoning, or integration across biological domains, Queryome consistently constructs explanations that trace why a conclusion follows from literature.

A central factor behind this behavior is Queryome’s multi-agent design. Instead of decomposing a question mechanically into subtasks, the principal investigator (PI) agent first performs a reconnaissance search to understand the structure of the problem. Only after identifying the relevant conceptual space does the PI decide whether to allocate planner–critic teams to pursue additional lines of inquiry. Queryome dynamically adjusts the depth and breadth of search. The architecture of Queryome is based on an important design principle grounded in recent findings about large language models: while LLMs perform small, well-scoped reasoning steps reliably, their accuracy drops sharply when asked to manage large, unfiltered context windows [27]. Queryome converts this limitation into an architectural advantage. Rather than overloading a single model with raw text, it distributes analysis into smaller, well-defined reasoning tasks under critic supervision. Each retrieved article is assessed, scored, and justified before contributing to synthesis, ensuring that every cited source plays a substantive role in the reasoning chain. We would also like to note that the performance of Queryome will continue to improve as more advanced LLMs become available, since these models can be directly incorporated as agents.

Despite these advances, Queryome remains constrained by its design choices. Limiting retrieval to PubMed ensures quality control but omits key knowledge sources such as clinical guidelines, preprints, and multimodal data. Its search policy, though effective, is still handcrafted rather than being data driven. Future directions include integrating reinforcement or meta-learning [54] to refine investigative heuristics, expanding to multilingual and multimodal retrieval [55], incorporating evidence quality weighting [56] based on study design and bias, and introducing temporal tracking [57] to monitor shifts in consensus. Equally important will be developing interfaces that make the system safe, transparent, and trustworthy for clinical or research use. Building upon this foundation, we plan to develop a more comprehensive version with access to a broader range of biological databases, wikis, and scientific literature. This larger system could also incorporate specialized agents capable of performing tasks like gene sequence similarity searches [58, 59] or incorporation of bioinformatics function [60, 61] and structure prediction [62–65] tools [66].

In the broader context, Queryome represents a step toward AI systems that engage directly with the empirical foundations of biomedicine. The long-term vision is not automation but collaboration: an ecosystem of reasoning agents that extend human scientific inquiry, preserving rigor while amplifying reach.

## Supporting information

Suppplementary information, tables, and a figure

## Acknowledgements

We thank Drs. Mark Christie, Angeline Lyon, and Janice Evans for expert evaluation of system outputs. This work was partly supported by the National Institutes of Health (R35GM158267, R21AI187928) and the National Science Foundation (IIS2211598, DBI2146026, and DBI2422620).

## Data and Code Availability

The source code of Queryome is available at https://github.com/kiharalab/queryome. In addition, an application that runs on MacOS (Intel and Apple Silicon) and Windows is available at: https://www.queryome.app/. Benchmark results can be downloaded from: https://kiharalab.org/queryome/benchmark_data.tar.gz.

## Author Contributions

PP conceived the study and developed Queryome. NI computed embeddings provided technical guidance. HS contributed to application development including database infrastructure. SG participated in the method design. ET participated in the expert evaluation. DK guided the benchmarking design, analyzed data. PP wrote the initial draft. NI and SG participated in writing the initial draft. Particularly, NI analyzed and wrote the Ablation studies section. DK critically edited it. All authors read and approved the manuscript.

