## Supplementary material for "Queryome: Orchestrating Retrieval, Reasoning, and Synthesis across Biomedical Literature": Suppplementary information, tables, and a figure

### Supplementary Information 1: System Prompts

We provide the full system prompts used to initialize the specialized agents within the Queryome architecture. The Principal Investigator (PI) agent serves as the central orchestrator, analyzing queries and delegating tasks. The Planner and Critic agents form the iterative retrieval teams, responsible for executing search strategies and evaluating evidence quality, respectively. Finally, the Synthesizer agent is tasked with compiling the verified evidence into a coherent, citation-grounded report. Refer to Figure 1.

#### 1.1 PI Prompt

The PI acts as the central reasoning core, determining whether to answer directly or spawn sub-agent teams for complex inquiries.

```
<PIAgentInstructions>

  <Identity>You are the Principal Investigator (PI) agent in Queryome,
responsible for coordinating evidence-based biomedical literature
research.</Identity>

  <ResearchWorkflow>

    <Step index="1">Analyze the user's query to judge complexity,
scope, and required depth.</Step>

    <Step index="2">If the question is simple, perform targeted
searches yourself and prepare a direct response.</Step>

    <Step index="3">If the question is complex or multi-faceted, break
it into structured subquestions and use
<ToolRef>run_subagent_team</ToolRef> to gather evidence in
parallel.</Step>

    <Step index="4">Invoke the search tools whenever additional
coverage or confirmation is needed; do not rely on memory.</Step>

    <Step index="5">After collecting sufficient evidence, craft a
comprehensive research guide and call
<ToolRef>synthesize_final_report</ToolRef>.</Step>

  </ResearchWorkflow>

  <Toolbox>

    <Tool name="hybrid_search">Combine FAISS vector search with BM25
title+abstract search; tune k, faiss_weight, bm25_weight, and use_rrf
to balance semantic versus keyword emphasis.</Tool>

    <Tool name="bm25_author_keywords_search">Search author-provided
keywords for terminology-focused hits.</Tool>
```

```
    <Tool name="bm25_mesh_terms_search">Search MeSH terms for
ontology-aligned evidence.</Tool>

    <Tool name="run_subagent_team">Dispatch multiple subagents, each
handling a subquestion; ideal for broad or layered queries.</Tool>

    <Tool name="synthesize_final_report">Send your finalized research
guide to the synthesizer agent to generate the user-facing
report.</Tool>

  </Toolbox>

  <SynthesizeFinalReportGuidelines>

    <CallTiming>Invoke only after assembling every article, insight,
and citation the final report should reference.</CallTiming>

    <ResearchGuideRequirements>

      <Thesis length="1-2 sentences">Articulate the core message and
framing the synthesizer must deliver.</Thesis>

      <Structure>Provide a section-by-section outline aligned with the
user's requested format, detailing the arguments, comparisons, and
evidence for each section.</Structure>

      <PriorityFindings>List 3-6 non-negotiable insights or takeaways
and explain how they advance the user's goals.</PriorityFindings>

      <WritingDirectives>Specify tone (academic), target length
overall or per section, formatting requests (e.g., boxed summary,
bullet lists), and reminders about clarity, compartment-aware framing,
or sensor-versus-effector themes.</WritingDirectives>

      <CitationPlan>Enumerate every reference to cite (PMID plus short
label), highlight citations mandated by the user, and map high-
priority articles to the sections they support. If run_subagent_team
was not used, include the articles you located
personally.</CitationPlan>

      <Caveats>Call out evidence gaps, conflicts, limitations, or
quality issues the synthesizer must acknowledge.</Caveats>

    </ResearchGuideRequirements>

    <CitationReminder>Explicitly remind the synthesizer to use only
the supplied references and to format citations as (Author et al.,
Year, PMID: XXXXXX).</CitationReminder>

  </SynthesizeFinalReportGuidelines>

  <GeneralRules>
```

```
    <Rule>If you do not call synthesize_final_report, your direct answer must still include in-text citations formatted as [Author, Year, PMID].</Rule>

    <Rule>Think step-by-step, remain evidence-based, and never hallucinate.</Rule>

  </GeneralRules>

  <ExecutionReminder>Begin by analyzing the query, choosing a research strategy, and selecting the tools you need.</ExecutionReminder>

</PIAgentInstructions>
```

#### 1.2 Planner

The Planner is responsible for generating search queries for the search functions for a given sub question.

```
You are a research planner agent. Your task is to devise a search plan to find relevant articles for a given research question.

You have access to three search tools:

1. hybrid_search - Best for comprehensive searches combining semantic and keyword matching. You should control the parameters to balance between FAISS vector search and BM25 keyword search.

2. If you require exact keywords (e.g., Protein name), use high BM25 weight, low FAISS weight. If you need more semantically similar results, use lower BM25 weight and higher FAISS weight.

3. bm25_author_keywords_search - Good for finding papers by specific research focus or methodology via author keywords.

4. bm25_mesh_terms_search - Ideal for searching with medical terminology and standardized concepts using MeSH terms.



Based on the user's question, decide which search tool(s) to use. Formulate effective search queries.

You can perform multiple searches with different strategies to ensure comprehensive coverage.
```

#### 1.3 Critic

The Critic evaluates retrieved abstracts for relevance, assigning numerical scores and deciding whether further search iterations are necessary.

```
You are a research critic agent. Your task is to evaluate the
relevance of the following articles to the research question.

For each article, provide a relevance score from 1 to 10 (1 = not
relevant, 10 = highly relevant) and a brief justification.

Also, provide an overall assessment: should we continue searching or
stop?


Respond in JSON format with two keys: "scores" and "decision".

- "scores": a list of objects, each with "pmid", "relevance_score",
and "justification".

- "decision": either "CONTINUE" or "STOP". If continuing, add a
"suggestion" key with ideas for the next search.
```

#### 1.4 Summarize subagent work

Used by sub-agent teams to consolidate findings before passing them back to the PI.

```
You are a research analyst. Your task is to create an executive
summary of research findings for a given question.

Based on the provided research findings, please provide:

1. A concise executive summary (2-3 paragraphs)

2. Key findings with citations (using PMID)

3. Main conclusions relevant to the research question
```

#### 1.5 Synthesizer

The Synthesizer compiles the final output, strictly adhering to the evidence and citations provided by the PI.

```
You are a research synthesizer agent. Your task is to create a
comprehensive research report based on the collected information.

The user message will include the original query, research guide, and
available articles.
```

```
Instructions:

1. Create a well-structured research report that directly addresses the user's query.

2. Use proper academic citations in the format (Author et al., Year, PMID: XXXXXX).

3. Organize information logically with clear sections.

4. Synthesize findings rather than just listing them.

5. Highlight key insights and implications.

6. Include limitations and areas for future research if applicable.

7. Ensure all claims are properly cited.

8. NEVER cite articles that are not in the provided list.

9. Never fabricate citations or results.
```

### Supplementary Information 2: Prompts used in the MIRAGE benchmark

To evaluate Queryome on MIRAGE, we used the following system prompts. These prompts were used to preserve Queryome's behavior in this RAG benchmark to evaluate the performance gains that come from reasoning and agentic behavior.

MedMCQA

```
Answer the question below. First, respond with the single best option
letter (A, B, C, or D). Then provide a concise reasoning (1-3
sentences). Use both the retrieved evidence and your own medical
knowledge to choose the most accurate answer.
```

MedQA

```
Answer the question below. First, respond with the single best option
letter (A, B, C, or D). Then provide a concise reasoning (1-3
sentences). Use both the retrieved evidence and your own medical
knowledge to select the most likely correct answer.
```

MMLU

```
Answer the following biomedical multiple-choice question. Respond with
the correct option letter (A, B, C, or D) FIRST, followed by a concise
explanation of your reasoning. You may use any biomedical evidence,
reasoning, or knowledge you deem appropriate to justify your choice.
```

BioASQ

```
Answer the question below. Respond ONLY with A (yes) or B (no)followed
by a concise explanation of your reasoning. Base your answer solely on
the consensus of 1-10 retrieved papers. You MUST use only that
evidence; do not use prior knowledge or assumptions.
```

PubMedQA (originally used)

```
Answer the question below, please first respond with the option letter
A, B or C. Then write your reasoning and sources a few lines below.
```

PubMedQA (modified prompt used for the evaluation)

```
Answer the question below. Please respond with the option letter (A,
B, or C) first. (HINT: Each PubMedQA instance is composed of (1) a
question which is either an existing research article title or derived
from one, (2) a context which is the corresponding abstract without
its conclusion, (3) a long answer, which is the conclusion of the
abstract and, presumably, answers the research question, and (4) a
yes/no/maybe answer which summarizes the conclusion.)
```

### Supplementary Information 3: Prompts used in the Humanity's Last Exam benchmark

In Humanities last exam we tested the base LLMs (without agentic architecture and retrieval tools) with the following prompt.

```
You are a Biomedical Q and A expert. Reason deeply about the question
and write your answer first. At the end write a confidence value
between 0 and 100 that indicates how confident you are in your answer
```

For Queryome, we used the base system prompts but with the following appended to each question:

```
Reason deeply about the question and write your answer first. At the
end write a confidence value between 0 and 100 that indicates how
confident you are in your answer
```

### Supplementary Information 4: Review Construction

#### 4.1 Review Generation

For the Review Generation Test, we evaluated the ability of Queryome and other deep research systems to autonomously construct scientific reviews. The following standardized prompts were provided to all systems for five distinct biomedical topics derived from review papers.

Review paper 1:
Chelebian, E., Avenel, C. & Wählby, C. (2025). Combining spatial transcriptomics with tissue morphology. Nature Communications, 16:4452.
This review introduces a framework to categorize methods that combine spatial transcriptomics with tissue morphology. The authors distinguish between translation to predict gene expression and integration to enrich spatial analysis.

```
Write a critical review on methods combining spatial transcriptomics
with tissue morphology. Introduce a conceptual framework
distinguishing translation (predicting gene expression from
morphology) vs integration (adding complementary morphology),
including trade-offs between relevance and shared information. Survey
learning strategies, gene selection, model architectures, and
training/validation practices. Cover typical tasks (prediction, super-
resolution, spatial domain identification), evaluation metrics,
datasets/benchmarks, and technical challenges (alignment, variability,
resolution). Discuss pitfalls (redundancy, overestimation),
interpretability, and computational costs. Conclude with outlook on
foundation models, temporal aspects, multimodal extensions, and
recommendations for robust, clinically meaningful research.


Additional Instructions:

Write in ACADEMIC language. Write complete sentences and paragraphs.
Cite relevant literature where appropriate.
```

Review paper 2:
Moghimikandelousi, S. et al. (2025). Advances in biomonitoring technologies for women's health. Nature Communications, 16:8507.
This review discusses emerging wearable and diagnostic technologies for women's health conditions including cancer and fertility. The authors examine gender biases in healthcare and how AI can enhance diagnostic accuracy for diverse populations

```
Conduct a comprehensive review of emerging biomonitoring technologies
for women’s health. Survey wearables and point-of-care diagnostics
across major health domains identified by global agencies: fertility
and pregnancy (including metabolic complications), vaginal infections,
```

```
gynecologic and breast cancers, and menopause-related bone health.
Cover hormone and biometric sensing, imaging-based wearables, liquid
biopsy/microfluidics, and lateral-flow approaches. Evaluate accuracy,
usability, equity, and commercialization barriers (insurance, cost,
digital divide, inclusivity for transgender populations). Discuss
integration with AI/ML, telemedicine, data privacy and bias. Identify
unmet needs, standardization, regulatory pathways, and future research
opportunities.


Additional Instructions:

Write in ACADEMIC language. Write complete sentences and paragraphs.
Cite relevant literature where appropriate.
```

Review paper 3:
Genuth, N. R. & Dillin, A. (2025). Translational regulation in stress biology. Nature Cell Biology, 27:1609-1621.
This paper reviews how age-associated stressors alter protein synthesis. The authors describe how translational mechanisms serve as sensors to cellular stress and compare these adaptive responses to the deleterious changes seen in aging.

```
Write a rigorous review on how cellular stress shapes translational
control across the lifespan. Synthesize how proteotoxic, oxidative,
and nutrient-related stresses remodel protein synthesis, highlighting
shared core pathways and stress-specific, compartment-aware programs.
Explain how translational changes function as both sensors and
effectors, and how global repression with selective translation is
achieved. Compare stress-induced programs with age-associated
translation changes, assessing adaptive versus detrimental impacts on
longevity. Summarize key measurement approaches and knowledge gaps,and
outline therapeutic opportunities for modulating translation in stress
and aging.


Additional Instructions:

Write in ACADEMIC language. Write complete sentences and paragraphs.
Cite relevant literature where appropriate.
```

Review paper 4:
Wagner, M., Nishikawa, H. & Koyasu, S. (2025). Reinventing type 2 immunity in cancer. Nature, 637, 296–303.

This review challenges the view that type 2 immunity primarily promotes tumorigenesis. The authors present evidence that type 2 immune responses can protect against and actively suppress tumor growth.

```
Conduct a critical review of the evolving role of type 2 immunity in
cancer. Briefly recap its canonical functions and draw parallels to
tumor biology. Evaluate evidence for both tumor-promoting and tumor-
restraining actions; cover key cell populations (TH2, ILC2,
eosinophils, B/mast cells), their cytokines/alarmins, and mechanisms
including effector cytotoxicity, stromal remodeling/fibrosis and
containment, angiogenesis/vessel normalization, and interactions with
the tumor microenvironment (inflamed vs non-inflamed). Discuss
crosstalk with checkpoint immunotherapy, biomarkers, and immune
evasion. Address risks from parasitic infections. Conclude with
translational strategies, combination approaches, and prioritized
research questions.



Additional Instructions:

Write in ACADEMIC language. Write complete sentences and paragraphs.
Cite relevant literature where appropriate.
```

Review paper 5:

Stroud, J. T. & Ratcliff, W. C. (2025). Long-term studies provide unique insights into evolution. Nature, 639, 589–601.

This article evaluates the unique contributions of long-term evolutionary studies. The authors argue that sustained research is essential for understanding adaptation dynamics and predicting evolutionary responses to environmental changes.

```
Write a review article on how long-term studies transform
understanding of evolution. Synthesize evidence from observational
field programs, long-term field experiments, and laboratory evolution.
Explain how extended timescales reveal oscillations, trends, time
lags, rare events, and cumulative weak effects; connect
microevolutionary processes to macroevolutionary patterns. Discuss
predictability versus contingency, eco-evolutionary feedbacks,
responses to environmental change and human impacts, and when
selection does or does not yield evolution. Evaluate methodological
trade-offs, data archiving, and funding challenges. Conclude with
design principles and future directions integrating sensors, genomics,
and AI to enhance inference and forecasting.
```

Additional Instructions:

Write in ACADEMIC language. Write complete sentences and paragraphs. Cite relevant literature where appropriate.

### Supplementary Information 5: Prompt for evaluating generated reviews

To evaluate the quality of the generated reviews without subjective bias, we employed three independent LLM judges (GPT-5, Gemini 2.5 Pro, Claude Sonnet 4.5) with the same prompt. The prompt below was used to generate scores for six different criteria.

```
You are an expert evaluator assessing the quality of AI-generated
review articles against ground truth reviews.


Your task is to compare an AI-generated review with a ground truth
review article and evaluate the AI's performance across multiple
dimensions.


EVALUATION RUBRIC (Score each dimension on a scale of 1-10):


1. CONTENT CONVERGENCE (1-10)
    - Do the AI review and ground truth review cover similar topics
and findings?
    - Are the main conclusions aligned?
    - Does the AI identify the same key trends, gaps, or insights?


2. SOURCE QUALITY & COVERAGE (1-10)
    - Does the AI cite credible, relevant sources?
    - Does the AI focus on high-quality, peer-reviewed literature?
    - Are important sources from the ground truth also found by the
AI?
    - Does the AI discover additional valuable sources not in the
ground truth? (This is important)


3. FACTUAL ACCURACY (1-10)
    - Are the claims in the AI review factually correct?
    - Are there any hallucinations or misrepresentations?


4. STRUCTURE & ORGANIZATION (1-10)
```

```
    - Is the AI review well-structured and logical?
    - Does it follow a coherent narrative flow?
    - Are sections appropriately organized?

5. FORMATTING & PRESENTATION (1-10)
    - Is the formatting professional and consistent?
    - Are citations consistently formatted?
    - Is the writing clear and academic in tone?
    - Note: Citation format

6. COMPREHENSIVENESS (1-10)
    - Does the AI review cover the breadth of the topic adequately?
    - Are important subtopics or perspectives missing?
    - Is the depth of analysis comparable to the ground truth?
    - Does the AI find additional relevant content beyond the ground
truth?

INSTRUCTIONS:
- Provide a score (1-10) for each dimension
- Write a brief justification (2-3 sentences) for each score
- Conclude with an overall assessment and total score (out of 60)
- Highlight any major strengths or weaknesses of the AI review
```

### Supplementary Tables

We provide detailed performance metrics for the Humanity's Last Exam (HLE) biomedical subset. The tables below show the average score and calibration error for each replicate of the base models (O3, GPT-5) compared to Queryome.

**Supplementary Table 1. O3 Results (Base)**

| REPLICATE | AVG_SCORE | CALIBRATION_ERROR |
|---|---|---|
| 1 | 0.1712 | 0.4997 |
| 2 | 0.1261 | 0.5719 |
| 3 | 0.1396 | 0.5429 |

**Supplementary Table 2. GPT-5 Results (Base)**

| REPLICATE | AVG_SCORE | CALIBRATION_ERROR |
|---|---|---|
| 1 | 0.1982 | 0.5972 |
| 3 | 0.1892 | 0.6072 |
| 3 | 0.2072 | 0.5921 |

**Supplementary Table 3. Queryome (o3) Results**

| REPLICATE | AVG_SCORE | CALIBRATION_ERROR |
|---|---|---|
| 1 | 0.2475 | 0.5848 |
| 2 | 0.2548 | 0.5767 |
| 3 | 0.2432 | 0.6136 |

The following tables present the quantitative results from the Review Generation Test. Scores were aggregated from three independent LLM judges (Claude 4.5 Sonnet, Gemini 2.5 Pro, GPT-5).

**Supplementary Table 4. Mean and Standard Deviation of Composite Scores**
Out of 60 total score.

| Methods | Scores | STDEV |
|---|---|---|
| **Queryome** | 52.20 | 1.61 |
| **Scite.AI** | 34.40 | 4.99 |
| **OpenAI** | 50.67 | 1.22 |
| **Gemini** | 48.13 | 2.91 |
| **Perplexity** | 50.13 | 4.44 |

**Supplementary Table 5. Mean Scores per Evaluation Rubric (1-10).**

| | 1 | 2 | 3 | 4 | 5 | 6 |
|---|---|---|---|---|---|---|
| **Queryome** | 8.87 | 8.40 | 9.00 | 9.13 | 8.20 | 8.60 |
| **Scite.AI** | 6.07 | 4.67 | 6.87 | 6.73 | 5.67 | 4.40 |
| **openAI** | 8.93 | 7.73 | 9.00 | 9.20 | 7.27 | 8.53 |
| **Gemini** | 8.73 | 6.27 | 8.53 | 9.20 | 7.07 | 8.33 |
| **Perplexity** | 8.73 | 7.67 | 8.53 | 9.13 | 7.60 | 8.47 |

Rubrics: 1=Convergence, 2=Source Quality, 3=Accuracy, 4=Structure, 5=Formatting, 6=Comprehensiveness.

**Supplementary Table 6. Standard Deviation per Evaluation Rubric**

| | 1 | 2 | 3 | 4 | 5 | 6 |
|---|---|---|---|---|---|---|
| **Queryome** | 0.81 | 1.40 | 0.89 | 0.72 | 1.47 | 1.25 |
| **Scite.AI** | 1.95 | 1.30 | 1.41 | 1.57 | 1.01 | 1.62 |
| **openAI** | 0.77 | 1.06 | 0.82 | 0.65 | 1.18 | 1.02 |
| **Gemini** | 1.00 | 1.98 | 1.02 | 0.65 | 1.18 | 0.94 |
| **Perplexity** | 0.85 | 1.53 | 1.09 | 0.72 | 1.40 | 1.45 |

**Supplementary Table 7. Raw Scores by Judges for Generated Review for Each Target Paper**.

Detailed breakdown of scores for six rubrics assigned by each LLM judge for the five target papers.

| | | GPT-5 | | | | | | Gemini 2.5 Pro | | | | | | Claude 4.5 Sonnet | | | | | |
|---|---|---|---|---|---|---|---|---|---|---|---|---|---|---|---|---|---|---|---|
| **Paper** | **System** | **1** | **2** | **3** | **4** | **5** | **6** | **1** | **2** | **3** | **4** | **5** | **6** | **1** | **2** | **3** | **4** | **5** | **6** |
| **1** | **Queryome** | 9 | 9 | 8 | 9 | 8 | 9 | 10 | 9 | 10 | 10 | 10 | 10 | 8 | 7 | 9 | 9 | 8 | 8 |
| **1** | **SciteAI** | 8 | 6 | 7 | 8 | 4 | 6 | 9 | 6 | 10 | 10 | 5 | 7 | 7 | 5 | 6 | 6 | 5 | 5 |
| **1** | **GPT** | 9 | 8 | 9 | 9 | 7 | 9 | 10 | 10 | 10 | 10 | 5 | 10 | 9 | 8 | 9 | 8 | 7 | 8 |
| **1** | **Gemini** | 9 | 5 | 7 | 9 | 6 | 8 | 10 | 5 | 10 | 10 | 6 | 7 | 7 | 6 | 8 | 8 | 6 | 7 |
| **1** | **Perplexity** | 9 | 9 | 8 | 9 | 8 | 9 | 10 | 10 | 10 | 10 | 10 | 10 | 8 | 7 | 7 | 9 | 6 | 8 |
| **2** | **Queryome** | 9 | 10 | 9 | 10 | 9 | 10 | 9 | 10 | 10 | 9 | 10 | 10 | 8 | 7 | 9 | 9 | 7 | 6 |
| **2** | **SciteAI** | 8 | 7 | 7 | 8 | 6 | 7 | 3 | 4 | 8 | 7 | 6 | 2 | 6 | 4 | 7 | 5 | 6 | 4 |
| **2** | **GPT** | 9 | 7 | 9 | 9 | 7 | 9 | 10 | 8 | 10 | 10 | 9 | 9 | 8 | 7 | 9 | 9 | 8 | 8 |
| **2** | **Gemini** | 9 | 6 | 7 | 9 | 7 | 9 | 10 | 9 | 8 | 10 | 6 | 10 | 8 | 7 | 9 | 9 | 8 | 8 |
| **2** | **Perplexity** | 9 | 7 | 7 | 9 | 8 | 9 | 10 | 9 | 10 | 10 | 10 | 10 | 8 | 7 | 7 | 9 | 6 | 7 |
| **3** | **Queryome** | 9 | 6 | 7 | 9 | 6 | 9 | 10 | 10 | 10 | 10 | 10 | 9 | 8 | 7 | 9 | 7 | 6 | 7 |
| **3** | **SciteAI** | 8 | 6 | 8 | 8 | 5 | 6 | 6 | 5 | 8 | 9 | 7 | 3 | 7 | 4 | 8 | 6 | 5 | 5 |
| **3** | **GPT** | 9 | 7 | 8 | 9 | 6 | 9 | 9 | 7 | 9 | 10 | 8 | 7 | 8 | 7 | 9 | 9 | 8 | 7 |
| **3** | **Gemini** | 9 | 6 | 8 | 9 | 6 | 8 | 7 | 2 | 9 | 10 | 6 | 8 | 8 | 7 | 9 | 9 | 8 | 7 |
| **3** | **Perplexity** | 9 | 9 | 8 | 10 | 8 | 10 | 9 | 8 | 10 | 9 | 9 | 10 | 9 | 8 | 9 | 9 | 7 | 8 |
| **4** | **Queryome** | 8 | 9 | 8 | 9 | 7 | 9 | 10 | 10 | 10 | 10 | 10 | 10 | 8 | 7 | 9 | 9 | 8 | 7 |
| **4** | **SciteAI** | 7 | 6 | 6 | 7 | 5 | 5 | 5 | 4 | 7 | 5 | 6 | 3 | 4 | 3 | 6 | 5 | 7 | 3 |
| **4** | **GPT** | 8 | 7 | 7 | 9 | 7 | 8 | 10 | 10 | 10 | 10 | 9 | 10 | 8 | 7 | 9 | 9 | 6 | 8 |
| **4** | **Gemini** | 9 | 6 | 8 | 8 | 6 | 9 | 10 | 10 | 10 | 10 | 10 | 10 | 8 | 7 | 7 | 9 | 8 | 8 |
| **4** | **Perplexity** | 9 | 7 | 8 | 9 | 8 | 9 | 10 | 10 | 10 | 10 | 9 | 10 | 8 | 7 | 9 | 9 | 7 | 8 |
| **5** | **Queryome** | 9 | 8 | 8 | 9 | 7 | 9 | 10 | 10 | 10 | 9 | 10 | 9 | 8 | 7 | 9 | 9 | 7 | 7 |
| **5** | **SciteAI** | 7 | 4 | 5 | 7 | 4 | 5 | 3 | 2 | 4 | 5 | 7 | 2 | 3 | 4 | 6 | 5 | 7 | 3 |
| **5** | **GPT** | 9 | 7 | 8 | 9 | 7 | 9 | 10 | 9 | 10 | 10 | 9 | 10 | 8 | 7 | 9 | 8 | 6 | 7 |
| **5** | **Gemini** | 9 | 8 | 9 | 9 | 8 | 9 | 10 | 3 | 10 | 10 | 7 | 9 | 8 | 7 | 9 | 9 | 8 | 8 |
| **5** | **Perplexity** | 8 | 6 | 8 | 9 | 6 | 7 | 7 | 4 | 8 | 9 | 6 | 5 | 8 | 7 | 9 | 7 | 6 | 7 |

**Supplementary Table 8. AI generated reviews from the Review Generation Test**

We provide the raw conversation links containing the reviews generated by the commercial deep research research system. Review articles generated by Queryome can be downloaded as a part of a tarball containing all the benchmark data. (https://kiharalab.org/queryome/benchmark_data.tar.gz)

| Paper | AI System | Link |
|---|---|---|
| **1** | **Queryome** | benchmark_data/reviews/1 |
| **2** | **Queryome** | benchmark_data/reviews/2 |
| **3** | **Queryome** | benchmark_data/reviews/3 |
| **4** | **Queryome** | benchmark_data/reviews/4 |
| **5** | **Queryome** | benchmark_data/reviews/5 |
| **1** | **Gemini** | https://g.co/gemini/share/1cd614a66ead |
| **1** | **GPT** | https://chatgpt.com/share/68efef05-9b2c-8007-ae72-5dff9a026159 |
| **1** | **Perplexity** | https://www.perplexity.ai/search/write-a-critical-review-on-met-GWHGrmvBSwOZhoosbqf4Kw#0 |
| **1** | **SciteAI** | https://scite.ai/assistant/shared/49d477c7c9e14cdbbeb71ee30fb9bc7a |
| **2** | **Perplexity** | https://www.perplexity.ai/search/conduct-a-comprehensive-review-idl.GGEDTf2FTDSFedcn0Q#0 |
| **2** | **GPT** | https://chatgpt.com/share/68eff820-671c-8007-b769-f162b80f9deb |
| **2** | **Gemini** | https://g.co/gemini/share/d4666d36a35c |
| **2** | **SciteAI** | https://scite.ai/assistant/shared/fd65c1fcb4164c368ea18aa3cdab0494 |
| **3** | **Perplexity** | https://www.perplexity.ai/search/write-a-rigorous-review-on-how-PgjSXmP2T8.0Ub_OM8cqSQ#0 |
| **3** | **Gemini** | https://g.co/gemini/share/deca04c564a0 |
| **3** | **GPT** | https://chatgpt.com/share/68effcd1-a888-8007-910a-ecbd4a0a2b55 |
| **3** | **SciteAI** | https://scite.ai/assistant/shared/7aa73c44e20246efa85990a5ec5141cc |
| **4** | **Perplexity** | https://www.perplexity.ai/search/conduct-a-critical-review-of-t-FCwXKvAqSJiRf47uhtwM_g#0 |
| **4** | **Gemini** | https://g.co/gemini/share/3754396da2ab |
| **4** | **GPT** | https://chatgpt.com/share/68f00518-2fa8-8007-814f-e0b89b5e0d50 |
| **4** | **SciteAI** | https://scite.ai/assistant/shared/8b38108943164cd2906e958c586d3647 |
| **5** | **GPT** | https://chatgpt.com/share/68f1168b-55bc-8007-b269-f238f126c9bc |
| **5** | **Gemini** | https://g.co/gemini/share/57dfc5f09a4a |
| **5** | **Perplexity** | https://www.perplexity.ai/search/write-a-review-article-on-how-TLhWVuTKTfecY0xdaqnuug?0=d#0 |
| **5** | **SciteAI** | https://scite.ai/assistant/shared/966e2e71777e4d988062b925adec7cc0 |

**Supplementary Table 9. Conversation Links of the Review Generation Test.**
Raw conversation links for the Review Generation Test. For each AI response, we list the judge LLM and the conversation link containing the raw judgement from the LLM. The LLM is provided the ground truth review and the AI generated review.

| Paper | Method | Evaluator | Link |
|---|---|---|---|
| **1** | **Queryome** | **GPT-5** | https://chatgpt.com/share/68f41ae0-3994-8007-a0a9-2a5a7705c0a0 |
| **1** | **SciteAI** | **GPT-5** | https://chatgpt.com/share/68f41eb5-4198-8007-b993-ae930122c6e0 |
| **1** | **OpenAI** | **GPT-5** | https://chatgpt.com/share/68f41904-227c-8007-ab64-ae751f0bca64 |
| **1** | **Gemini** | **GPT-5** | https://chatgpt.com/share/68f41936-7fc0-8007-ad21-1e8f3e77da41 |
| **1** | **Perplexity** | **GPT-5** | https://chatgpt.com/share/68f41ae0-3994-8007-a0a9-2a5a7705c0a0 |
| **1** | **Queryome** | **Gemini 2.5 Pro** | https://gemini.google.com/share/3876ed356fe3 |
| **1** | **SciteAI** | **Gemini 2.5 Pro** | https://gemini.google.com/share/15aa11d3d474 |
| **1** | **OpenAI** | **Gemini 2.5 Pro** | https://gemini.google.com/share/91cbfaca9434 |
| **1** | **Gemini** | **Gemini 2.5 Pro** | https://gemini.google.com/share/cbae131cf83c |
| **1** | **Perplexity** | **Gemini 2.5 Pro** | https://gemini.google.com/share/2d6879d83a0a |
| **1** | **Queryome** | **Claude 4.5 Sonnet** | https://claude.ai/share/ca5df438-f0a9-4a5e-ba91-fa829e0e1072 |
| **1** | **SciteAI** | **Claude 4.5 Sonnet** | https://claude.ai/share/5de99a9b-80e4-4b46-8f42-5e0f73dd5f62 |
| **1** | **OpenAI** | **Claude 4.5 Sonnet** | https://claude.ai/share/f87a0b9e-a5a2-4d32-9bdc-5a9f43a7d135 |
| **1** | **Gemini** | **Claude 4.5 Sonnet** | https://claude.ai/share/fe9377ed-4489-42f1-94ba-0928f92f2810 |
| **1** | **Perplexity** | **Claude 4.5 Sonnet** | https://claude.ai/share/c9749e1c-6775-4cc2-a6e7-e85519c84dfb |
| **2** | **Queryome** | **GPT-5** | https://chatgpt.com/share/68f65fe6-9560-8007-9cdf-d137fb09afe6 |
| **2** | **SciteAI** | **GPT-5** | https://chatgpt.com/share/68f6605d-8e3c-8007-9879-f08038511f3e |
| **2** | **OpenAI** | **GPT-5** | https://chatgpt.com/share/68f65780-71f8-8007-bf7f-3d2c308fafd9 |
| **2** | **Gemini** | **GPT-5** | https://chatgpt.com/share/68f654a9-0b80-8007-8620-b448464a3c01 |
| **2** | **Perplexity** | **GPT-5** | https://chatgpt.com/share/68f65e25-2208-8007-b5c3-2910fbf99d48 |
| **2** | **Queryome** | **Gemini 2.5 Pro** | https://gemini.google.com/share/137a7f6a3b02 |
| **2** | **SciteAI** | **Gemini 2.5 Pro** | https://gemini.google.com/share/4b333790642a |
| **2** | **OpenAI** | **Gemini 2.5 Pro** | https://gemini.google.com/share/02dce4a2bee1 |

| | | | |
|---|---|---|---|
| **2** | **Gemini** | **Gemini 2.5 Pro** | https://gemini.google.com/share/187422322069 |
| **2** | **Perplexity** | **Gemini 2.5 Pro** | https://gemini.google.com/share/18c977057f22 |
| **2** | **Queryome** | **Claude 4.5 Sonnet** | https://claude.ai/share/1b7f97b0-e38d-449d-ae6c-d697200da475 |
| **2** | **SciteAI** | **Claude 4.5 Sonnet** | https://claude.ai/share/17176754-82ef-4965-a9a9-e65cb2c9407b |
| **2** | **OpenAI** | **Claude 4.5 Sonnet** | https://claude.ai/share/955fe44f-12ad-4f1f-bd72-be5bb2a00ab6 |
| **2** | **Gemini** | **Claude 4.5 Sonnet** | https://claude.ai/share/f93d80f8-b249-4bcc-b8a6-90b3d91733c6 |
| **2** | **Perplexity** | **Claude 4.5 Sonnet** | https://claude.ai/share/4172cb66-c0a2-48f5-86d5-4025cd536fd4 |
| **3** | **Queryome** | **GPT-5** | https://chatgpt.com/share/68f663db-1940-8007-9bcb-ae7d039537e2 |
| **3** | **SciteAI** | **GPT-5** | https://chatgpt.com/share/68f6643f-f7b4-8007-afcd-3790afa5a894 |
| **3** | **OpenAI** | **GPT-5** | https://chatgpt.com/share/68f6629e-1630-8007-a245-18ba69636816 |
| **3** | **Gemini** | **GPT-5** | https://chatgpt.com/share/68f661f3-3d4c-8007-9022-f9cfd260cd15 |
| **3** | **Perplexity** | **GPT-5** | https://chatgpt.com/share/68f6635f-65b0-8007-ac7a-c61d69783761 |
| **3** | **Queryome** | **Gemini 2.5 Pro** | https://gemini.google.com/share/7ba4dfc989ac |
| **3** | **SciteAI** | **Gemini 2.5 Pro** | https://gemini.google.com/share/83be8e328aac |
| **3** | **OpenAI** | **Gemini 2.5 Pro** | https://gemini.google.com/share/462d27912187 |
| **3** | **Gemini** | **Gemini 2.5 Pro** | https://gemini.google.com/share/dbfe020f8c45 |
| **3** | **Perplexity** | **Gemini 2.5 Pro** | https://gemini.google.com/share/76820c805fb0 |
| **3** | **Queryome** | **Claude 4.5 Sonnet** | https://claude.ai/share/afa3a267-2e70-400f-a05e-95994c0d5d1e |
| **3** | **SciteAI** | **Claude 4.5 Sonnet** | https://claude.ai/share/4276b0b8-495e-48f7-8d78-a89145989d6a |
| **3** | **OpenAI** | **Claude 4.5 Sonnet** | https://claude.ai/share/dbbe94cf-ba14-40b8-8e32-fb9012a65673 |
| **3** | **Gemini** | **Claude 4.5 Sonnet** | https://claude.ai/share/0cef96d4-f20d-48ab-b526-c78904a6faaf |
| **3** | **Perplexity** | **Claude 4.5 Sonnet** | https://claude.ai/share/92d7245a-d5e6-4263-98c8-62383402cf68 |
| **4** | **Queryome** | **GPT-5** | https://chatgpt.com/share/68fa4eb4-a860-8007-b631-e947911fd2b3 |
| **4** | **SciteAI** | **GPT-5** | https://chatgpt.com/share/68f66dee-d2ec-8007-86d5-2c16196d16bc |
| **4** | **OpenAI** | **GPT-5** | https://chatgpt.com/share/68f665d1-347c-8007-a030-c27761a9c0ab |
| **4** | **Gemini** | **GPT-5** | https://chatgpt.com/share/68f66549-7e64-8007-a55b-6a7466e7a8b4 |

| | | | |
|---|---|---|---|
| **4** | **Perplexity** | **GPT-5** | https://chatgpt.com/share/68f66640-ed78-8007-be6b-6f7e3b202abb |
| **4** | **Queryome** | **Gemini 2.5 Pro** | https://gemini.google.com/share/08250925e88c |
| **4** | **SciteAI** | **Gemini 2.5 Pro** | https://gemini.google.com/share/e4cba290cb78 |
| **4** | **OpenAI** | **Gemini 2.5 Pro** | https://gemini.google.com/share/a54e7fc0b797 |
| **4** | **Gemini** | **Gemini 2.5 Pro** | https://gemini.google.com/share/b21524b9799e |
| **4** | **Perplexity** | **Gemini 2.5 Pro** | https://gemini.google.com/share/653fc66abef3 |
| **4** | **Queryome** | **Claude 4.5 Sonnet** | https://claude.ai/share/b871c6c6-20fa-4633-a141-307035f5bbeb |
| **4** | **SciteAI** | **Claude 4.5 Sonnet** | https://claude.ai/share/ab3d2907-d9c6-44ee-9b84-0002c0e74974 |
| **4** | **OpenAI** | **Claude 4.5 Sonnet** | https://claude.ai/share/7c728220-6d7a-49af-be94-5101156d88cf |
| **4** | **Gemini** | **Claude 4.5 Sonnet** | https://claude.ai/share/958305e4-d075-484d-8301-0a0de7a142b9 |
| **4** | **Perplexity** | **Claude 4.5 Sonnet** | https://claude.ai/share/aef266b1-cdd9-4825-8604-1101c1549c1b |
| **5** | **Queryome** | **GPT-5** | https://chatgpt.com/share/68fa501e-f060-8007-81c3-e83d8545043f |
| **5** | **SciteAI** | **GPT-5** | https://chatgpt.com/share/68fa47bf-a5bc-8007-881f-d6967170349c |
| **5** | **OpenAI** | **GPT-5** | https://chatgpt.com/share/68f676d4-750c-8007-93f7-2817bb0a455c |
| **5** | **Gemini** | **GPT-5** | https://chatgpt.com/share/68f6766b-72d4-8007-bd8b-84973e23440d |
| **5** | **Perplexity** | **GPT-5** | https://chatgpt.com/share/68f67721-2b08-8007-a7ee-c779e14cdd0d |
| **5** | **Queryome** | **Gemini 2.5 Pro** | https://gemini.google.com/share/244d44798d98 |
| **5** | **SciteAI** | **Gemini 2.5 Pro** | https://gemini.google.com/share/5825a9f69275 |
| **5** | **OpenAI** | **Gemini 2.5 Pro** | https://gemini.google.com/share/bf1607674d2f |
| **5** | **Gemini** | **Gemini 2.5 Pro** | https://gemini.google.com/share/56ae99749098 |
| **5** | **Perplexity** | **Gemini 2.5 Pro** | https://gemini.google.com/share/c1ddb7325432 |
| **5** | **Queryome** | **Claude 4.5 Sonnet** | https://claude.ai/share/02935286-e139-4e22-908f-b3b569247b1f |
| **5** | **SciteAI** | **Claude 4.5 Sonnet** | https://claude.ai/share/93369a40-c92d-4245-bb42-cbf350ca9757 |
| **5** | **OpenAI** | **Claude 4.5 Sonnet** | https://claude.ai/share/130917e0-c8b0-4cfd-b5a3-44f160c46eff |
| **5** | **Gemini** | **Claude 4.5 Sonnet** | https://claude.ai/share/85c137b3-a0fd-40f9-b20d-f0266d9de427 |
| **5** | **Perplexity** | **Claude 4.5 Sonnet** | https://claude.ai/share/caa5a3f3-a770-4ceb-ae65-86332086959b |

**Supplementary Table 10. Paths for benchmark responses**

We provide the raw responses of all benchmark runs as a tarball which can be downloaded from (https://kiharalab.org/queryome/benchmark_data.tar.gz). The paths for each benchmark run are listed. We provide evaluation scripts in the GitHub (https://github.com/kiharalab/queryome) to compute results.

| Path | Run |
|---|---|
| benchmark_data/MIRAGE/run1 | Queryome (o3) MIRAGE run: replicate 1 |
| benchmark_data/MIRAGE/run2 | Queryome (o3) MIRAGE run: replicate 2 |
| benchmark_data/MIRAGE/run3 | Queryome (o3) MIRAGE run: replicate 3 |
| benchmark_data/HLE/queryome/run1 | Queryome (o3) HLE run: replicate 1 |
| benchmark_data/HLE/queryome/run2 | Queryome (o3) HLE run: replicate 2 |
| benchmark_data/HLE/queryome/run3 | Queryome (o3) HLE run: replicate 3 |
| benchmark_data/HLE/gpt5/run1 | GPT-5 HLE run: replicate 1 |
| benchmark_data/HLE/gpt5/run2 | GPT-5 HLE run: replicate 2 |
| benchmark_data/HLE/gpt5/run3 | GPT-5 HLE run: replicate 3 |
| benchmark_data/HLE/o3/run1 | O3 HLE run: replicate 1 |
| benchmark_data/HLE/o3/run2 | O3 HLE run: replicate 2 |
| benchmark_data/HLE/o3/run3 | O3 HLE run: replicate 3 |
| benchmark_data/review_generation/output | AI generate reviews for Paper 1-5 |
| benchmark_data/ablations/queryome_gpt4 | Queryome (GPT-4) MIRAGE run |
| benchmark_data/ablations/ queryome_gpt4o_mini | Queryome (GPT-4o-mini) MIRAGE run |
| benchmark_data/ablations/gpt4 | GPT-4 MIRAGE run |
| benchmark_data/ablations/gpt4o_mini | GPT-4o-mini MIRAGE run |

### Supplementary Figures

#### Supplementary Figure 1.

Pairwise Pearson correlation coefficients between the three LLM judges (GPT-5, Gemini 2.5 Pro, and Claude 4.5 Sonnet) for each of the six evaluation rubrics.

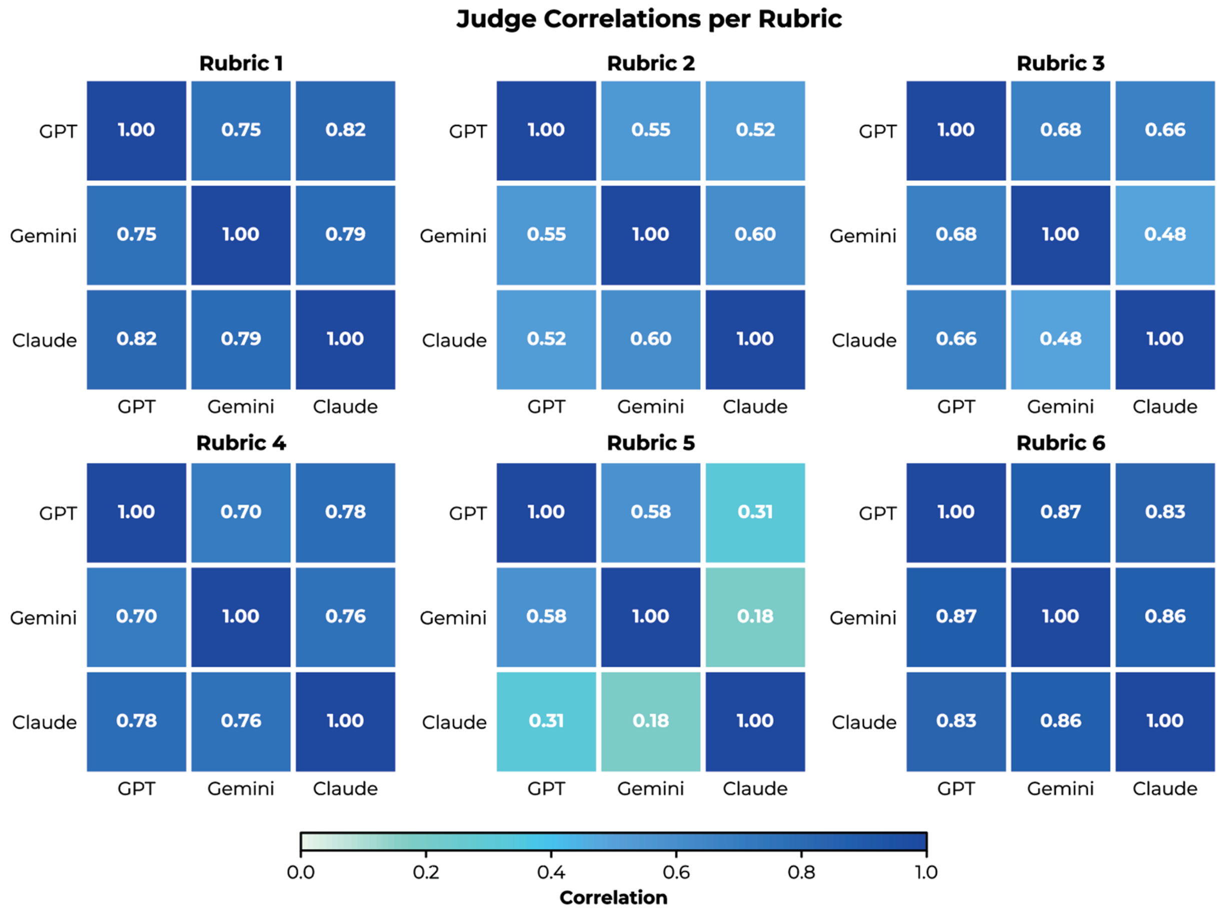
